# Prominent involvement of acetylcholine in shaping stable olfactory representation across the *Drosophila* brain

**DOI:** 10.1101/2024.04.03.587915

**Authors:** Jiaqi Fan, Yuling Wang, Lingbo Li, Jing He, Zhifeng Zhao, Fei Deng, Guochuan Li, Xinyang Li, Yiliang Zhou, Jiayin Zhao, Yulong Li, Jiamin Wu, Lu Fang, Qionghai Dai

## Abstract

Despite the vital role of neuromodulation in the neural system, the specific spatiotemporal dynamics of neuromodulators and their interactions with neuronal activities *in vivo* are still unclear, hampering our understanding of their information representation and functional contributions systemically. To address this problem, we employed two-photon synthetic aperture microscopy (2pSAM) to record long-term neuronal and neuromodulatory olfactory responses across the *Drosophila* brain at high speed. Our results revealed distinct response properties, global information propagation, functional connectivity, and odor identity representation among neuronal, cholinergic, and serotoninergic dynamics across multiple brain regions. We discovered the compensation between neuronal activity and cholinergic dynamics, both in the odor identity representation across the brain and the functional connectivity network structures of specific brain regions. Moreover, employing low-dimensional manifold and functional connectivity network analyses, we characterized the stable representation of cholinergic dynamics over a long term. Collectively, our unbiased and comprehensive investigation unveiled the prominent involvement of acetylcholine (ACh) in shaping olfactory representation across the brain, underscoring the inadequacy of solely considering neuronal activities when examining information representation of the brain.

## Main

Neuromodulation is a pervasive and pivotal feature of the neural system, exerting its influence on multiple aspects and scales, such as modulating attention and brain states^1,2^, shaping neuronal circuits^3,4^, and sensory processing^5,6^. However, the spatiotemporal dynamics of neuromodulators across the brain remain unclear, hindering our understanding of their functional roles systemically^7–9^. For instance, in the context of sensory processing, while different neuromodulators have distinct cellular effects, the purported functions overlap substantially^10,11^. Thus the current hypotheses of the functions and roles of particular neuromodulators may be limited^7^. Furthermore, the relationship between neuronal activity and neuromodulator release *in vivo* is poorly characterized due to limited indicators and imaging systems for simultaneous recording^8,12–16^. Consequently, how neuronal and neuromodulatory activities interact and synergistically represent and process information remains largely unknown. Previous research reported dissociable dopamine dynamics, indicating possible firing-independent dopamine release and highlighting the insufficiency of solely recording dopamine cells for understanding dopamine signals^15^. Therefore, elucidating the spatiotemporal dynamics and information representation of neuronal and neuromodulatory activities in a systemic view is critical for advancing our understanding of the neural system.

A typical example is the representation of sensory information, specifically olfaction, which is considered a fundamental cognitive function of the brain with significant involvement of neuromodulation^5,6,17,18^. Despite the longstanding research interest in olfactory processing, comprehension of olfactory representation in terms of neuronal activity and neuromodulator dynamics remains largely unexplored. The current understanding of olfactory processing is biased towards early brain layers, leaving olfactory information representation and arrangement in higher-order regions elusive^19–25^. Moreover, intriguingly, there is evidence of representational drift in neuronal activity related to olfaction and other sensory modalities occurring over hours and days^26,27^, yet the extent of these changes requires further characterization. Neuromodulators, such as acetylcholine (ACh)^11,28–30^ and serotonin (5-HT)^6,7,10^, play crucial roles in olfactory processing and memory. However, their spatiotemporal dynamics in relation to olfactory stimuli and the collective representation of olfactory information by neuronal and neuromodulatory activities across the brain over extended periods remain open questions. Recent rapid advancements in fluorescent neuromodulator indicators and high-throughput microscopy techniques offer the potential for unbiased large field-of-view (FOV) simultaneous recording of neuronal and neuromodulatory activities, which can enlighten us regarding these unresolved issues. A recent study revealed spatiotemporal heterogeneous coordination of cholinergic and neocortical activity across different waking states through simultaneous wide-field recording^8^. There is still a gap in understanding information representation by neuronal and neuromodulatory activities at high resolution.

Here, we employed two-photon synthetic aperture microscopy (2pSAM) for long-term high-speed multiple-brain-region volumetric imaging^31^ of the *Drosophila* brain, which was pan-neuronally labeled by the green-fluorescent calcium indicator jGCaMP7f (G7f)^32^ and a red-fluorescent indicator (rGRAB_ACh-0.5 (rACh) for ACh, rGRAB_HTR2C-0.5 (r5-HT) for 5-HT)^33,34^, to elucidate odor representation by neuronal and neuromodulatory activities across the *Drosophila* brain. Unsupervised deep-learning-based denoising algorithms were applied to increase the signal-to-noise ratio^35–37^. Our findings revealed diverse response properties, global information propagation, functional connectivity, and odor representation across the brain and among the indicators (G7f, rACh, and r5-HT). Despite extensive cholinergic connections across the *Drosophila* brain^38,39^, odor responses and discrimination of ACh were particularly significant in the Mushroom body (MB), especially the medial lobe (MBML), and adjacent brain regions, in contrast to the widespread coding observed in neuronal activity. Whereas the multiple-brain-region odor identity representation abilities of calcium and ACh were similar. Furthermore, integrating ACh dynamics improved the performance of odor identity representation by calcium, implying additional information beyond direct correlation with neuronal activity. Notably, voxel-level functional connectivity networks of ACh in specific brain regions^40^ also exhibited compensation and accentuation to the connectivity structures of neuronal activity. By analyzing the low-dimensional manifolds of olfactory responses, we further characterized odor-specific and neurochemical-specific representations and the temporal changes. We discovered that ACh dynamics exhibited superior stability compared to neuronal activities or 5-HT dynamics in the low-dimensional manifolds and functional connectivity networks over a long period. These findings may open up a new horizon for the systemic study of information representation and the neural networks encompassing neuronal and neuromodulatory dynamics, and uncover the essential role of neuromodulators in information representation.

### Multiple-brain-region recording of neuronal and neuromodulatory activities

To simultaneously record neuronal and neuromodulatory activities across the *Drosophila* brain, we employed a three-step crossbreeding method to generate flies with pan-neuronal expression of G7f together with either rACh or r5-HT. We surgically exposed the whole central brain by gently opening the posterior head cuticle of the fly and keeping the posterior brain surface flat. 2pSAM was used for dual-color volumetric imaging of the *Drosophila* brain at 30 Hz (Fig. 1a), with the FOV measuring 458.7 μm × 458.7 μm × 100 μm. This FOV covered nearly the entire lateral range and about half of the axial range of the fly central brain (about 80 to 180 μm under the surface), corresponding to approximately 43 brain regions (Fig. 1a-c; Supplementary Table 1 and Supplementary Video 1)^41,42^.

**Fig. 1.**
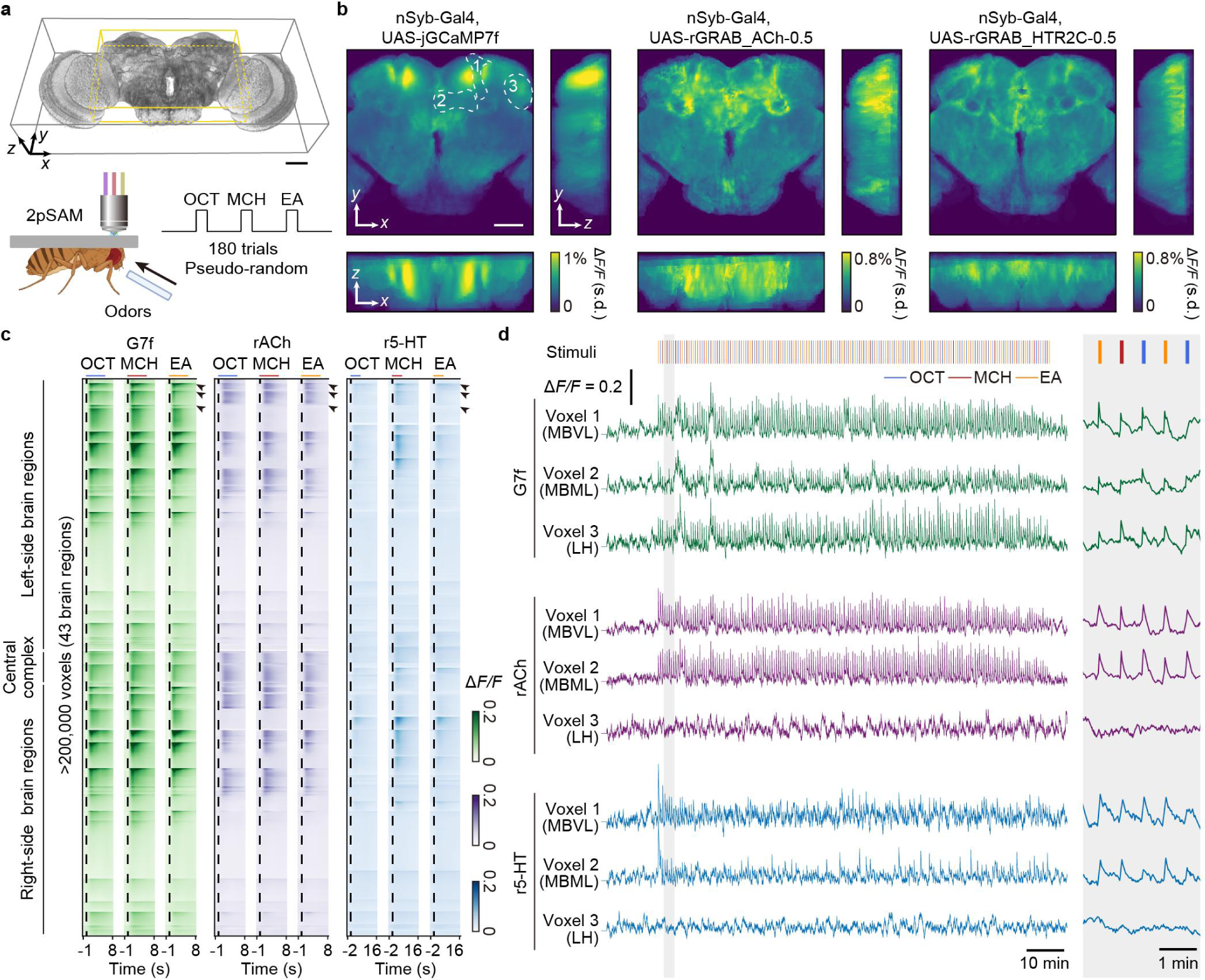
Long-term high-speed volumetric imaging of neuronal and neuromodulatory activities across the *Drosophila* central brain by 2pSAM. **a,** Experimental set-up and FOV in the *Drosophila* brain. Bottom: Imaging set-up (left) and experimental paradigm (right). A mounted fly is placed under the objective, exposing the posterior of the brain to light. 2pSAM performs low-phototoxicity fast 3D imaging by needle-like beam scanning. Three odors (OCT, MCH, EA) with a pseudo-random order are delivered to the antennas of the fly in 180 trials to test the long-term odor responses. Top: FOV in the *Drosophila* brain shown in 3D schematic diagram. The FOV covers nearly the whole lateral range and about half of the axial range of the fly central brain (about 80 to 180 μm under the surface, including about 43 brain regions, labeled by the yellow box). **b**, Average odor response intensities of three indicators, G7f, rACh, and r5-HT, across the FOV. 10 flies co-labeled by G7f and rACh and 10 flies co-labeled by G7f and r5-HT are analyzed. *n* = 20 flies for G7f, *n* = 10 flies for rACh, *n* = 10 flies for r5-HT. Flies co-labeled by G7f and rACh with high-intensity non-specific fluorescence on the upper edge of the brain are excluded for clarity. **c**, Trial-averaged G7f, rACh, and r5-HT responses to 3 odor stimuli of every voxel in the FOV of a fly. More than 200,000 voxels in 43 brain regions throughout the central brain are recorded simultaneously at 30 Hz. The voxels are arranged in the order of brain regions and the voxels within a brain region are listed by the response intensity from high to low. Since r5-HT response is significantly slower, its response in 16 s after odor delivery is shown, while 8 s is shown for G7f and rACh. Odor stimulus periods (5 s) are shown as lines above (blue: OCT; red: MCH; orange: EA). **d**, Left: Example Δ*F/F* traces of the voxels in 3 brain regions in **c** (labeled by black arrowheads). Right: The zoomed-in traces of the labeled period (gray shade). The vertical short lines above sign odor stimulus periods (blue: OCT; red: MCH; orange: EA). Heterogeneous spatiotemporal response between indicators can be observed. Regions MBVL, MBML, and LH of the right semi-brain are labeled by dashed lines with numbers 1, 2, and 3 in **b**. G7f and rACh responses are from a co-labeled fly, and the r5-HT response is from another fly co-labeled by G7f and r5-HT (the G7f response of this fly is not shown for concision). Scale bars: 50 μm in **a**, **b**.

To study neuronal and neuromodulatory activities during odor stimulation, we administered three different single compounds as olfactory stimuli—3-octanol (OCT), 4-methylcyclohexanol (MCH), and ethyl acetate (EA)—to the flies in a random order (Fig. 1a and Supplementary Video 1). At the concentration adopted in this study, OCT and MCH are two comparably representative aversive odors for flies, whereas EA is relatively more attractive^43–46^, which enable the investigation of odor identity as well as latent preference. Utilizing the 3-dimensional (3D) imaging ability and low phototoxicity of 2pSAM, we conducted long-term volumetric recording of neuronal and neuromodulatory activities at a sampling rate of 30 Hz for approximately two hours, with 180 trials (60 sessions) of odor stimulation in total, preceded by a 10-minute resting-state period (Fig. 1a, d). This approach facilitated the analysis of odor representation stability and temporal changes brain-widely. Throughout the imaging process, the flies maintained a favorable condition and exhibited consistent odor responses (Fig. 1d). We employed high-performance denoising algorithms^35–37^ to extract the temporal traces with a high signal-to-noise ratio (SNR) for more than 200,000 voxels across multiple brain regions per fly (Fig. 1c and Extended Data Fig. 1a). Notably, apparent heterogeneity and distinction of responses are observed for calcium, ACh, and 5-HT (Fig. 1b-d; and Supplementary Video 1), which will be discussed in detail in the following section.

### Diverse olfactory responses and global information propagation across the brain

Calcium, ACh and 5-HT showed significant responses to odor stimuli. To demonstrate the spatiotemporal dynamics of these neurochemicals, we analyzed the distribution, intensity, and dynamic characteristics of their responses and observed diverse response patterns and global information propagation across the brain.

To analyze the distribution of responses, we calculated the correlation between odor stimulus and trial-averaged Δ*F/F* as a measurement of responsiveness. We discovered that, for all three indicators, most brain regions showed high average correlation (Fig. 2a and Extended Data Fig. 1b), indicating a widespread response. However, 5-HT demonstrated a relatively lower responsiveness. At the brain-region level, the correlation in MB was relatively low for G7f but high for rACh; what’s more, correlations in the Superior lateral protocerebrum (SLP) and Lateral horn (LH) were high for G7f, but low for rACh and r5-HT, respectively (Fig. 2a and Extended Data Fig. 1b).

**Fig. 2.**
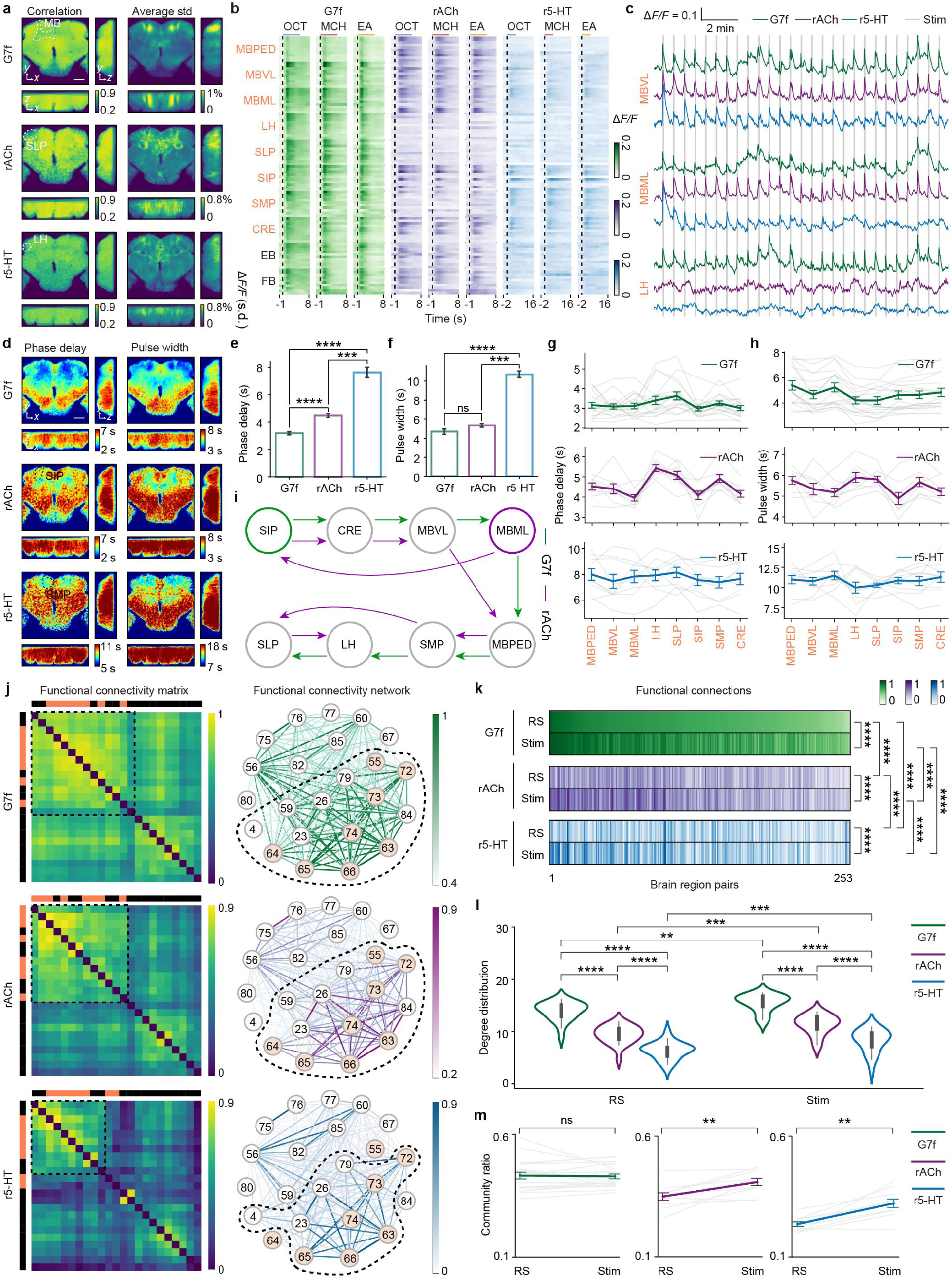
Heterogeneous and distinct olfactory responses and functional connectivity across the brain for G7f, rACh, and r5-HT. **a**, Left: Maps of correlations between trial-averaged G7f, rACh, or r5-HT activities and odor stimulus, averaged across flies. Right: Maps of average standard deviations of Δ*F/F* under odor stimuli across trials, averaged across flies. The dashed lines label the specific brain regions. **b**, Trial-averaged G7f, rACh, and r5-HT responses to 3 odor stimuli of the sample voxels in 8 olfactory regions (coral) and 2 non-olfactory regions (black) of a fly. Dashed lines sign the start of odor delivery. Short lines above sign odor stimulus periods (5 s) (blue: OCT; red: MCH; orange: EA). G7f and rACh responses are from a co-labeled fly, and the r5-HT response is from another fly co-labeled by G7f and r5-HT. **c**, Δ*F/F* traces of some voxels in MBVL, MBML, and LH in several trials. The gray lines sign the start of odor delivery. **d**, Maps of phase delays (left) and pulse widths (right) of odor responses for G7f, rACh, and r5-HT, averaged across flies. The dashed lines label the specific brain regions. **e**, **f**, Statistics of the average phase delay (**e**) and pulse width (**f**) of olfactory regions in the left semi-brain among G7f, rACh, and r5-HT. **g**, **h**, Statistics of phase delay (**g**) and pulse width (**h**) of each indicator in each olfactory brain region in the left semi-brain. **i**, Schematic of the global olfactory information propagation across olfactory regions for G7f (green) and rACh (purple) inferred from the phase delay. **j**, Left: Clustered functional connectivity matrices during odor stimulation of the left-side brain regions and the central complex for each indicator. The dashed boxes sign the communities containing most olfactory regions. The color blocks above and to the left of each matrix mark 8 olfactory (coral) and 15 non-olfactory regions (black). Right: Functional connectivity networks based on the matrices. The colors and widths of the edges indicate connection strengths, and the colors of the nodes indicate olfactory (coral) or non-olfactory (blank) regions. The dashed circles sign the communities in the dashed boxes on the left. The numbers labeled on the nodes indicate brain region indices, corresponding to the region names in Extended Data Fig. 3a. For the functional connectivity of the resting state, see Extended Data Fig. 3b. **k**, The functional connection patterns among three indicators during odor stimulation and the resting state. **l**, Statistics of the degree distribution in functional connectivity networks among three indicators during odor stimulation and the resting state. **m**, Statistics of the community ratio of each indicator during odor stimulation and the resting state. 10 flies co-labeled by G7f and rACh and 10 flies co-labeled by G7f and r5-HT are analyzed. *n* = 20 flies for G7f, *n* = 10 flies for rACh, *n* = 10 flies for r5-HT, mean ± s.e.m. A fly co-labeled by G7f and rACh with an obvious lean angle of the brain is excluded for clarity in (**a**, left). Flies co-labeled by G7f and rACh with high-intensity non-specific fluorescence on the upper edge of the brain are excluded for clarity in (**a**, right). A fly co-labeled by G7f and rACh with the interference of the calculation of phase delay by the non-specific fluorescence on the upper edge of the brain is excluded in **d-h**. Flies without a general coverage of region LH are excluded in **g**, **h**. Each light-colored line represents the result of a fly in **g**, **h**, and **m**. Stim: Odor stimulation. RS: The resting state. Two-sided Mann-Whitney U test in **e**, **f**; Scheirer–Ray–Hare test (non-parametric two-way ANOVA test) in **k**; Kolmogorov-Smirnov test in **l**; Two-sided Wilcoxon signed-rank test in **m**; \*\*\*\**P* < 0.0001, \*\*\**P* < 0.001, \*\**P* < 0.01, \**P* < 0.05, ns - not significant (*P* > 0.05). Scale bars: 50 μm in **a**, **d**.

The response intensity exhibited larger heterogeneity, with only a small number of voxels demonstrating very high values (Fig. 2a, Methods). Consequently, when assessing the average intensity of every brain region, no significant elevation was observed in the olfactory brain regions compared to other regions (Extended Data Fig. 1c). Notably, we noticed low intensity in LH and SLP for rACh and r5-HT contrasting with G7f (Fig. 2a-c). This finding is surprising considering that the lateral protocerebrum is recognized as a crucial higher-level olfactory center, where neuromodulation is deemed vital^22^. In contrast, the relative intensity in MB was high for rACh, surpassing even that of G7f (Fig. 2b, c), implying a potentially superior role of ACh in MB^39,47^. At the voxel level, there was substantial variation in response intensity even within a single brain region (Figs. 1c and 2b). Interestingly, the response intensities of G7f and rACh exhibited some degree of complementarity, with specific voxels displaying low intensity for G7f but high intensity for rACh (Fig. 2b, c).

To assess the dynamic properties, we calculated the phase delays and pulse widths of the odor responses for each indicator as measurements of response delay and duration (Extended Data Fig. 2a, Methods). The overall phase delay of G7f across multiple brain regions was significantly lower compared to the other indicators (Fig. 2d, e; and Extended Data Fig. 2b-d), identifying a broad quick neuronal response. The phase delay of MB was lower than SLP and LH (Fig. 2d, g). In contrast to the broad quick response of calcium, the phase delay of rACh was larger, with MBML and the Superior intermediate protocerebrum (SIP) showing apparent quicker response among the olfactory regions (Fig. 2d, e, and g; and Extended Data Fig. 2b). The phase delay of r5-HT was the largest, with slightly quicker response mainly in the Superior medial protocerebrum (SMP), SIP, and the Vertical lobe of mushroom body (MBVL) (Fig. 2d, e, and g; and Extended Data Fig. 2b). The average pulse widths of G7f and rACh were comparable and significantly lower than r5-HT, indicating the longer influence of r5-HT (Fig. 2d, f; and Extended Data Fig. 2e). The distributions of pulse width and phase delay across brain regions were similar for rACh. Larger phase delay came with the larger pulse width. However, the distributions displayed approximately opposite trends for G7f and r5-HT (Fig. 2d, g, and h). Notably, heterogeneity was observed within each region when analyzing phase delay and pulse width distributions of all voxels in the olfactory regions (Extended Data Fig. 2c, d, f, and g). Overall, the response dynamics and global information propagations differed among G7f, rACh, and r5-HT. We depicted a schematic illustrating the global information propagations for G7f and rACh based on the phase delay of each olfactory region (Fig. 2i). The result for r5-HT was not included due to high individual differences in phase delay (Fig. 2g).

To further characterize the distinction of calcium, ACh, and 5-HT dynamics at the system level, we constructed functional connectivity matrices and networks for each indicator based on the activities during odor stimulation (Fig. 2j and Extended Data Fig. 3a) and the resting state (Extended Data Fig. 3b). In general, G7f displayed high global connectivity, while rACh and r5-HT showed higher contrast in the connections. The networks exhibited distinct connectivity patterns across indicators and states, with some connections remaining consistent (Fig. 2j, k; and Extended Data Fig. 3a, b). During odor stimulation, two primary communities were observed in the networks (Methods), with similar community divisions among the three indicators. Whereas, LH and the Pedunculus of mushroom body (MBPED) were separated from other olfactory regions in the r5-HT communities (Fig. 2j). G7f displayed higher node degree compared to rACh and r5-HT in both states (Fig. 2l and Extended Data Fig. 3c). The connection strength ratio of the major olfactory-related community (community ratio) for rACh was comparable to G7f and higher than r5-HT during odor stimulation, suggesting strong olfactory-related connections for ACh and calcium and weaker connections for 5-HT (Extended Data Fig. 3d). Examining the network changes in relation to odor stimulation compared with the resting state, all indicators exhibited altered degree distributions across the states, with the average node degrees increasing for G7f and r5-HT while remaining stable for rACh (Fig. 2l; and Extended Data Fig. 3e, f). The community ratio of G7f remained consistent across states, indicating a stable olfactory-related community significance under both conditions (Fig. 2m; and Extended Data Fig. 3a, b). However, significant changes were observed in this ratio for rACh and r5-HT, suggesting notable connectivity alterations in response to odor stimulation (Fig. 2m; and Extended Data Fig. 3a, b).

### Odor identity representation by calcium, ACh, and 5-HT dynamics across multiple brain regions

We observed strong voxel-level response variations among different odors (Fig. 2b). To further examine odor identity representation, we conducted odor identity classification based on neuronal and neuromodulatory responses. Initially, we computed the classification accuracy of each block with a size of 4 × 4 × 2 voxels and generated accuracy maps across the FOV for each indicator (Fig. 3a). These accuracy maps revealed distinct spatial distributions of odor representations. G7f demonstrated widespread odor coding in multiple brain regions, particularly with high accuracy in LH (Fig. 3a, b). However, SLP and LH displayed low decoding accuracy for both rACh and r5-HT (Fig. 3b and Extended Data Fig. 1d), consistent with their low response levels (Fig. 2a-c; and Extended Data Fig. 1b, c). rACh showed high decoding accuracy predominantly in MB, consistent with its strong cholinergic response (Fig. 2a-c; and Extended Data Fig. 1b, c), indicating the superior role of ACh in MB. While olfactory regions did not exhibit significantly higher responsiveness or response intensity compared to other regions for all three indicators, most olfactory regions surpassed non-olfactory regions in accuracy (Fig. 3b and Extended Data Fig. 1).

**Fig. 3.**
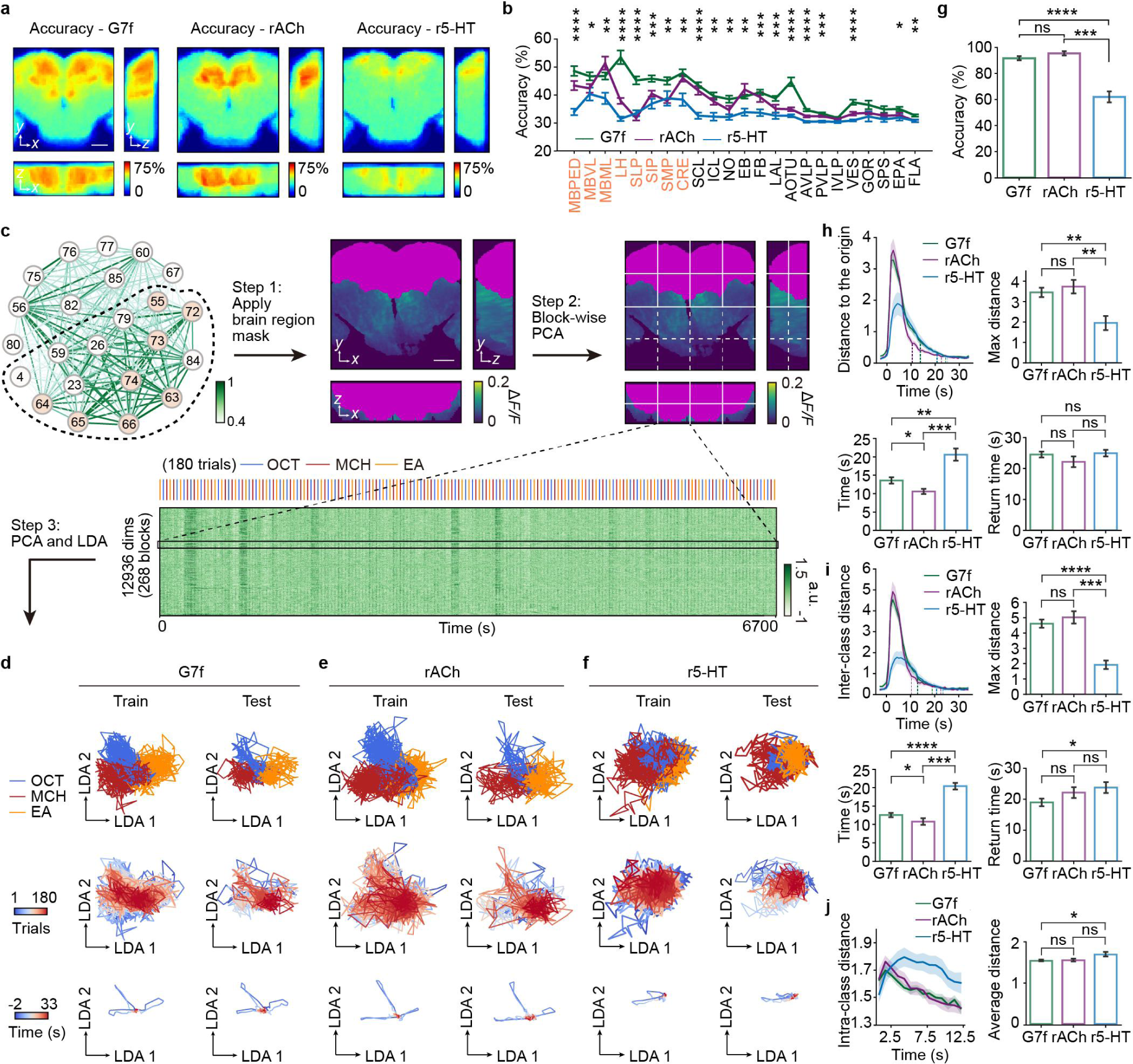
Odor identity representation by G7f, rACh, and r5-HT across multiple brain regions. **a**, Decoding accuracy maps of odor identity classification throughout the FOV, averaged across flies, for G7f, rACh, and r5-HT, from left to right. **b**, Statistics of the average decoding accuracy in each brain region for G7f, rACh, and r5-HT. Hypothesis testing is performed among three indicators in each brain region. Results of the left-side brain regions and the central complex are shown. **c**, Schematic of the dimensionality reduction method for voxel-level multiple-brain-region odor identity classification. Step 1: A brain region mask is applied according to the functional connectivity network to select the voxels within the olfactory-region-containing community. Step 2: Block-wise PCA efficiently reduces the data dimensionality, and the PCs from all blocks are consolidated. Step 3: PCA and LDA are implemented to further reduce the dimensionality and distinguish the odor identities within a low-dimensional space. **d-f**, Low-dimensional manifolds of the G7f (**d**), rACh (**e**), and r5-HT (**f**) responses to odor stimuli of a fly (left column: training set; right column: testing set). Top and middle: Each line is the odor response of a trial. Bottom: Each line is the average odor response of an odor identity. Manifolds from 2s before odor delivery to 33s after are plotted. Top: Colors denote odor identities (blue: OCT; red: MCH; orange: EA). Middle: Colors denote the trial numbers (blue: early; red: late). Bottom: Colors denote the time relative to odor delivery (blue: early; red: late). The manifolds in **d** and **e** are from the same fly. The arrow lines are arbitrary units but indicate an equivalent length in each dimension. **g**, Statistics of the voxel-level multiple-brain-region odor identity classification accuracies by G7f, rACh, and r5-HT. **h**-**j**, Metrics of the manifolds for three indicators. **h**, Top left: Average distance to the origin changing with time relative to odor delivery. The opaque dashed lines sign the time for returning to the one-fifth of the maximum distance. The high-transparency dashed lines sign the the time for returning to the random state. Top right: Statistics of the maximum distance. Bottom left: Statistics of the time for returning to the one-fifth of the maximum distance. Bottom right: Statistics of the time for returning to the random state. **i**, Similar to **h**, but for inter-class distance. **j**, Left: Average intra-class distance changing with time relative to odor delivery. Right: Statistics of the average intra-class distance within the period shown in the left figure. 10 flies co-labeled by G7f and rACh and 10 flies co-labeled by G7f and r5-HT are analyzed. *n* = 20 flies for G7f, *n* = 10 flies for rACh, *n* = 10 flies for r5-HT, mean ± s.e.m. Kruskal-Wallis test in **b**; Two-sided Mann-Whitney U test in **g**-**j**; \*\*\*\**P* < 0.0001, \*\*\**P* < 0.001, \*\**P* < 0.01, \**P* < 0.05, ns - not significant (*P* > 0.05). Scale bars: 50 μm in **a**, **c**.

Importantly, we noticed heterogeneity in response properties and accuracies within single brain regions. We then compared the decoding accuracies of the average response of a brain region and the dimensionality reduction result of the multi-voxel responses within the same region to show the necessity for high-resolution recording (Methods). This analysis specifically focused on the olfactory regions (Methods; and Extended Data Fig. 4a, b). We found significant elevation in accuracy (Extended Data Fig. 5a-c), indicating the coordinated representation of odor identity within individual brain regions. We also compared the accuracies of two co-labeled indicators (G7f and rACh, or G7f and r5-HT) in various regions. In most regions, the accuracy of r5-HT was either lower or comparable to that of G7f. In contrast, rACh outperformed G7f in specific brain regions, such as MBML and EB (Extended Data Fig. 5d).

Furthermore, we employed dimensionality reduction to explore the representation of odor identity across multiple brain regions (Fig. 3c; Methods; and Extended Data Fig. 4c-f). Firstly, as the brain regions were divided into two communities according to the functional connectivity (Fig. 2j), we applied a brain-region mask to select the voxels within the community containing olfactory regions. Secondly, we utilized block-wise principal component analysis (PCA) to extract the principal components (PC) from each block, effectively reducing data dimensionality and ensuring computational efficiency. Thirdly, we consolidated the PCs from all blocks and implemented PCA again, followed by linear discriminant analysis (LDA), to map the high-dimensional activity into a stimulus-related low-dimensional space. LDA was performed on the training sets, and the resultant transformation was applied to the testing sets without any overlap in trials to avoid data leakage (Fig. 3d-f; and Supplementary Video 1). In the low-dimensional space, individual trial traces formed a manifold, with distinct trajectories separating trials associated with different odor stimuli and resulting in segregation. The trial order did not exhibit discernible separation, demonstrating representation stability. Trials originated from a random state around the center point, extended during odor presentation, and then gradually returned back. Subsequently, we applied a support vector machine (SVM) for odor identity classification in the LDA space. Instead of classifying the trials solely based on the responses at a specific time point^17,48^, we leveraged the whole traces during a period in the low dimensional space, considering the availability of more comprehensive information. Accuracies were assessed through 5-fold cross-validation. The voxel-level multiple-brain-region data yielded significantly higher accuracy than region-level data (Extended Data Figs. 4g, h, and 5e-g) and single brain regions (Extended Data Fig. 5h-j), indicating the coordinated olfactory representation across the brain. The overall performances for G7f and rACh were comparable (accuracies above 90%), while the accuracy for r5-HT was significantly lower (accuracy around 60%, Fig. 3g), as depicted in the manifolds (Fig. 3d-f).

Flies exhibited diverse motions during the experimental process. As it has been validated that extensive neuronal activities are related to motions and behaviors^42,49–51^, here comes a question of whether motions interfere with our analyses of odor identity representation. We captured videos of fly abdomens and extracted the motions (Extended Data Fig. 6a). The correlation between odor stimuli and motion energy varied a lot among different flies, showing weak, positive, or negative associations (Extended Data Fig. 6b). Behavioral features were extracted through PCA (Extended Data Fig. 6c), capable of predicting stimulus periods and intervals to some extent but unable to classify odor identities (Extended Data Fig. 6d). We further predicted the PCs of the neuronal and neuromodulatory activities from behavior, stimulus, and both (Extended Data Fig. 6e-m). Both behavior and stimulus could explain the partial variance of the activities. Intriguingly, behavior accounted for variance in rACh and r5-HT dynamics more than G7f (Extended Data Fig. 6f, i, and l). However, the activities explained by behavior could not discriminate odor identities, and removing the components explained by behavior did not affect odor identity classification (Extended Data Fig. 6g, j, and m). Therefore, we conclude that motion modulates neuronal and neuromodulatory activities but does not account for odor identity representation in our analysis.

### Low-dimensional manifolds uncover the characteristics of odor representation

The low-dimensional manifolds provide additional insights into the characteristics of odor representation (Fig. 3d-f). We measured some indices for quantitative analyses. During odor presentation, the manifolds expand and subsequently contract, resulting in an increase and subsequent decrease in the distance to the origin (Fig. 3h). G7f and rACh show significantly larger maximum distances compared to r5-HT, indicating greater dynamic changes in the low-dimensional space (Fig. 3h). However, the time taken to return to the random state is similar for all indicators (Fig. 3h). We also measured the inter-class (Fig. 3i) and intra-class distances (Fig. 3j) of these indicators. r5-HT represents a lower inter-class and higher intra-class distance, suggesting the poorer ability to distinguish odor identities. G7f demonstrates a slightly faster return to the undistinguishable state compared to r5-HT (Fig. 3i). Comparing the time from odor delivery to returning to the one-fifth of the maximum distance, rACh returns the fastest, and G7f returns faster than r5-HT (Fig. 3h, i). Overall, G7f and rACh perform similarly in the low-dimensional manifolds and outperform r5-HT.

The manifolds of three odors, OCT, MCH, and EA, also exhibit distinct characteristics. In the G7f manifolds (Extended Data Fig. 7a), EA (attractive odor) has a shorter maximum distance to the origin and a faster return than OCT and MCH (aversive ones). However, such distinctions are absent in the manifolds of rACh and r5-HT. OCT and MCH (O-M, both negative) are harder to classify than other odor pairs for G7f and rACh, as the maximum inter-class distance of O-M is apparently lower and the recovering time is significantly shorter, especially for G7f (Extended Data Fig. 7b). No significant differences are observed in the intra-class distance across odors (Extended Data Fig. 7c). The evident disparate manifestation of EA and the relatively low discrimination of O-M might be relevant to odor preference^43–46^, which is an aspect of odor perception better uncovered in higher-order brain regions^23,52,53^ and is likely to be reflected in our multiple-brain-region analysis. This effect is less evident for rACh and r5-HT than G7f.

### Integration of ACh and calcium dynamics improves the odor identity representation

Given that rACh and r5-HT respond to odor stimuli and represent odor identities in distinct manners from G7f, the questions arose: Does the information from these indicators complement that of neuronal activity? Can they enhance odor identity representation? We explored these questions and came to the conclusion that integrating ACh rather than 5-HT dynamics improves odor identity representation by neuronal activity.

By integrating the signals of rACh or r5-HT with G7f, we could conduct odor identity classification utilizing the dual-channel data (Methods). Initially, we observed significant accuracy increase in many individual brain regions in the accuracy map when integrating rACh signals (Fig. 4a, b). This was further validated by the voxel-level classification within each region (Extended Data Fig. 5d). Subsequently, we integrated the rACh signals from the brain regions with accuracy gain to the multiple-brain-region responses of G7f and performed the voxel-level multiple-brain-region odor identity classification. Remarkably, we observed substantial accuracy improvement upon integrating rACh signals at the multiple-brain-region level (Fig. 4c). This improvement was also evident in the manifolds (Fig. 4d, e; and Supplementary Video 1), where odor responses extended farther than either single indicator and trials with different odor identities separated more distinctly (Fig. 4f-h). For comparisons, integrating 5-HT did not yield similar accuracy gains (Extended Data Figs. 5d and 8, and Supplementary Video 1).

**Fig. 4.**
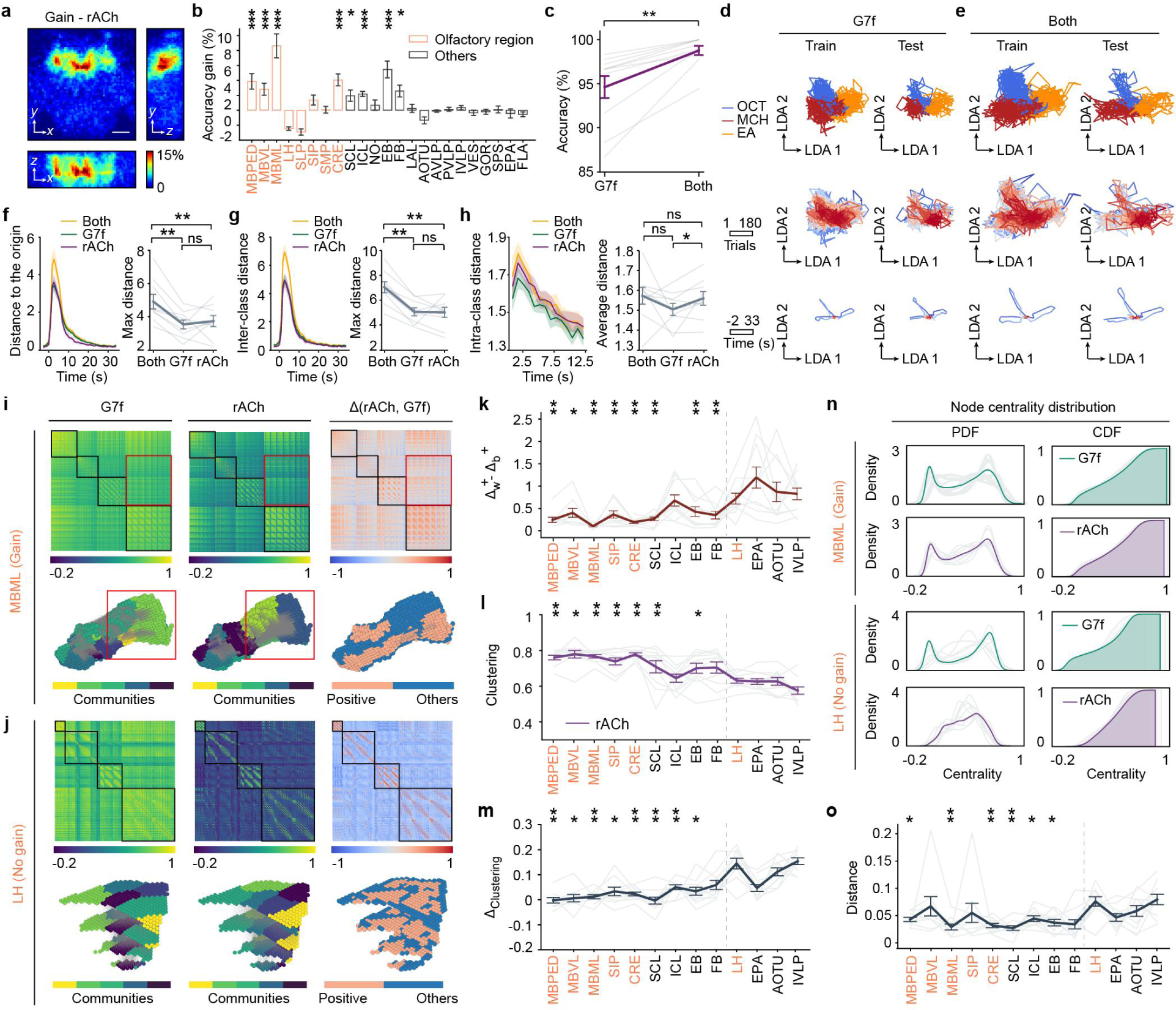
Integration of ACh dynamics improves the odor identity representation by neuronal activity. **a**, Map of decoding-accuracy gain by integrating rACh dynamics, averaged across flies. **b**, Statistics of the average accuracy gain in each brain region. **c**, Comparison of the voxel-level multiple-brain-region odor identity classification accuracies between using only the G7f channel and integrating both channels. **d**, **e**, Low-dimensional manifolds by G7f (**d**) and both channels (**e**) of a fly, similar to Fig. 3d. The manifolds are from the same fly as Fig. 3d. The arrow lines are arbitrary units but indicate an equivalent length in each dimension. **f-h**, Metrics of the manifolds for integrating both channels, G7f only and rACh only. **f**, Distance to the origin. **g**, Inter-class distance. **h**, Intra-class distance. **i**, Functional connectivity matrices (top) and mappings of communities and connectivity change in the physical space (bottom) of the voxels within MBML. Three columns are descriptions of the functional connectivity of G7f, rACh, and the difference in the deflation ratio between them, from left to right. There is connectivity compensation (top, red rectangles) and emphasis (top, black rectangles) to G7f by rACh. The compensation is reflected in the different distribution of functional connections in the physical space (bottom, red rectangles). The community divisions are similar for the two channels (bottom, labeled in different colors), and voxels with the greatest increased connectivity gather in space (bottom, coral). **j**, Similar to **i**, but for LH, a brain region without accuracy gain. Connectivity emphasis exists (black rectangles) in this region without compensation. **k**, The difference in increased deflation ratios within and between clusters of the functional connectivity of rACh compared to G7f. **l**, The average clustering coefficient of each brain region for rACh. **m**, The average clustering coefficient difference between rACh and G7f in each region. **n**, The node centrality distribution of MBML and LH. **o**, The distance of the node centrality distributions between rACh and G7f in each region. In **k**, **l**, **n**, and **o**, the brain regions on the left side of the dashed line are regions with accuracy gain, and the right ones are regions without gain; hypothesis testing is performed for each brain region with accuracy gain toward the average level of LH, EPA, AOTU, and IVLP. *n* = 10 flies co-labeled by G7f and rACh, mean ± s.e.m. Results of the left-side brain regions and the central complex are shown in **b**, **i-o**. Each light-colored line represents the result of a fly in **c**, **f-h**, and **k-o**. One-sided Wilcoxon signed-rank test in **b**, **l**, **m**, **o**; Two-sided Wilcoxon signed-rank test in **c**, **f-h**, **k**; \*\*\*\**P* < 0.0001, \*\*\**P* < 0.001, \*\**P* < 0.01, \**P* < 0.05, ns - not significant (*P* > 0.05). Scale bar: 50 μm in **a**.

Further, we examined the voxel-level functional connectivity networks within specific brain regions to investigate the neural mechanisms underlying the accuracy gain. Our goal was to identify differences between regions with and without accuracy gain and determine the possible source of the observed gain. Surprisingly, we discovered clear evidence of connectivity compensation and emphasis to G7f by rACh in brain regions with gain (Fig. 4i, k; and Extended Data Fig. 9a). Conversely, regions without gain only displayed connectivity emphasis without any noticeable compensation (Fig. 4j, k; and Extended Data Fig. 9a). In regions with accuracy gain, the community divisions (Methods) of the two channels exhibited similarities, with voxels demonstrating increased connectivity clustering together in space (Fig. 4i and Extended Data Fig. 9a). However, in regions without accuracy gain, although the community divisions were similar for the two channels, voxels with increased connectivity were not spatially clustered (Fig. 4j and Extended Data Fig. 9a). These phenomena were absent in the networks of the resting state (Extended Data Fig. 9b). Brain regions with gain also displayed stronger clustering and higher node heterogeneity than other regions for rACh, along with more similar clustering level and node centrality distribution to G7f (Fig. 4l-o; and Extended Data Fig. 9c). These results suggest that while the functional connectivity networks of the two indicators in brain regions with accuracy gain show apparent compensation, their network properties are similar, indicating strong functional segregation and rich information coding for both channels^54^. However, in regions without gain (e.g., LH), rACh displayed lower clustering and a more concentrated distribution of node centrality, indicating larger node homogeneity and lower functional segregation (Fig. 4l-o; and Extended Data Fig. 9c). The functional connectivity networks of r5- HT showed similar characteristics (Extended Data Fig. 8). The connectivity compensation while maintaining the network characteristics shows a local mismatch of ACh release and neuronal activity and should relate to the accuracy gain.

### ACh exhibits better temporal stability than calcium and 5-HT in odor representation

Since we found ACh is involved in odor identity representation with compensation to neuronal activities, we further investigated the temporal stability of ACh during odor representation. With the long-term low-phototoxicity imaging capability of 2pSAM, we successfully recorded consistent odor responses in 180 trials (60 sessions) of each fly. We evenly partitioned the 60 sessions into four stages (S1, S2, S3, S4) and obtained the low-dimensional manifolds for every stage (Fig. 5a). Analyzing the properties of the low-dimensional manifolds, we observed that G7f consistently exhibited a gradual decrease in the maximum distance to the origin and inter-class distance across stages, indicating a continuous reduction in representational amplitude and odor discrimination (Fig. 5b, c). In contrast, rACh remained stable, at least during the first three stages, while the reduction in r5-HT occurred primarily between S1 and S2. Intriguingly, G7f exhibited a progressive shift in the returning location of sessions, particularly from S1 to S2, suggesting a potential internal state change (Fig. 5a, d; Extended Data Fig. 10a, b). However, this change was not observed for rACh and r5-HT. Notably, the average intra-class distances of all indicators exhibited a significant decrease from S1 to S2, potentially indicating a process of learning or habituation (Fig. 5e).

**Fig. 5.**
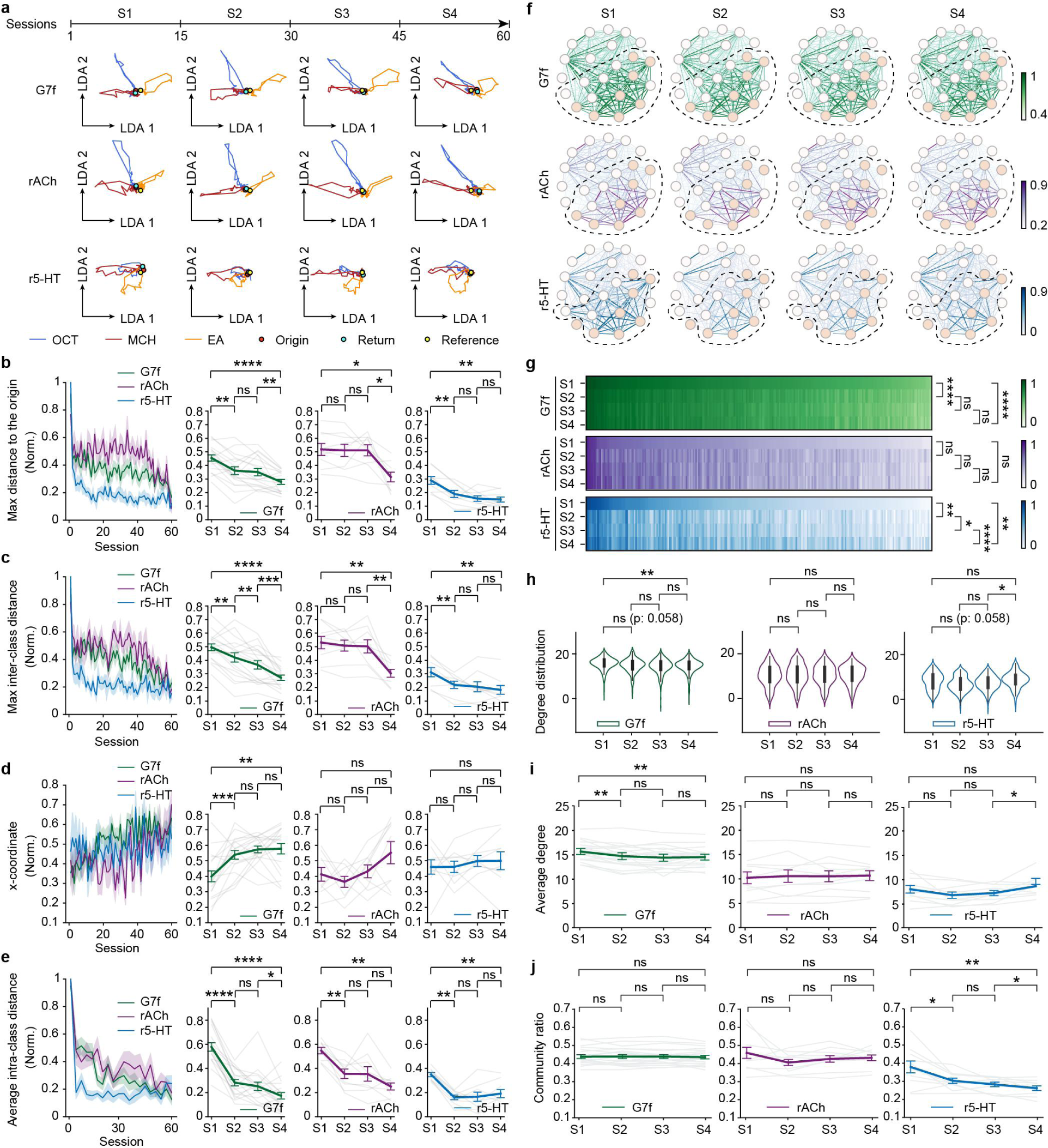
ACh exhibits better temporal stability in odor representation and the functional connectivity network than calcium and 5-HT. **a**, Average manifolds of a fly across 4 stages. S1: Sessions 1-15; S2: Sessions 16-30; S3: Sessions 31-45; S4: Sessions 46-60. Each session includes 3 trials. The arrow lines are arbitrary units but indicate an equivalent length in each dimension. Colors denote odor identities (blue: OCT; red: MCH; orange: EA). The red, cyan, and yellow dots sign the origins of the manifolds, the return locations of the manifolds, and the reference center points, respectively. **b**-**e**, Metrics of the manifold change across four stages. **b**, Maximum distance to the origin. **c**, Maximum inter-class distance. **d**, The x-coordinate of the return locations. **e**, Average intra-class distance. Left: Changes of the metrics across sessions. Right: Statistics of the metrics across four stages for each indicator. **f**, Functional connectivity networks of the brain regions for each indicator during odor stimulation in four stages. Consistent with Fig. 2j, the colors and widths of the edges indicate connection strengths, and the colors of the nodes indicate olfactory (coral) and non-olfactory (blank) regions. The dashed circles sign communities in Fig. 2j. **g**, The functional connection patterns of three indicators among four stages. **h**, Statistics of the degree distribution in functional connectivity networks of three indicators among four stages. **i**, The statistics of the average degree of the functional connectivity networks among four stages. **j**, Statistics of the community ratio in the functional connectivity networks of three indicators among four stages. 10 flies co-labeled by G7f and rACh and 10 flies co-labeled by G7f and r5-HT are analyzed. *n* = 20 flies for G7f, *n* = 10 flies for rACh, *n* = 10 flies for r5-HT, mean ± s.e.m. Each light-colored line represents a fly in **b**-**e**, **i**. Two-sided Wilcoxon signed-rank test in **b**-**e**, **i**; Scheirer–Ray–Hare test (non-parametric two-way ANOVA test) in **g**; Kolmogorov-Smirnov test in **h**; \*\*\*\**P* < 0.0001, \*\*\**P* < 0.001, \*\**P* < 0.01, \**P* < 0.05, ns - not significant (*P* > 0.05).

Overall, rACh exhibited the most stable odor representation across stages, while G7f displayed a continuous and gradual change. r5-HT demonstrated significant reductions from S1 to S2, consistent with the dramatic decrease in response happening only in the several trials at the very beginning (Figs. 1d and 2c). We further characterized the functional connectivity networks of the three indicators, assessing functional connection patterns, degree distribution, and community ratio across the four stages (Fig. 5f-j; and Extended Data Fig. 10c). The results corroborated our findings from the analysis of the low-dimensional manifolds. While the functional network of ACh is very stable across 2 hours, the functional networks of calcium and 5-HT both change within 2 hours. All these results indicate the stable odor representation of ACh.

## Discussion

High-throughput neuronal recording techniques have brought many discoveries either in the spatial or the temporal domain^26,27,42,55–61^, inaccessible for localized studies. Understanding the spatiotemporal dynamics and information representation of neuronal activity serves as a fundamental step for a deeper and more comprehensive understanding of functions. In contrast, large-scale neuromodulator dynamics still lack studying despite their vital role in the neural system, due to technical limitations^8^. Therefore, abundant mysteries might be hidden in neuromodulator dynamics as their quantitative relationship with neuronal activity *in vivo* remains elusive^8,12,13,15^. Our current understanding of neuromodulatory functions, primarily derived from targeted investigations, might be limited^7^.

The latest advancement in high-performance fluorescent neuromodulator indicators^33,34,62^ and microscopic techniques^31,35–37,63^ has enabled us to investigate the spatiotemporal dynamics of neuromodulators and neuronal activity, explore their involvement in information representation functionally, and examine the relationship between neuronal and neuromodulatory activities (Fig. 1). To address these objectives, we focused on olfactory processing as a context. Initially, we identified the diverse response properties and functional connectivity across neuronal, cholinergic, and serotoninergic dynamics (Fig. 2). Furthermore, we investigated the odor identity representation across multiple brain regions, identifying the low-dimensional manifolds and the latent compensation of information to neuronal activity by ACh (Figs. 3 and 4). Notably, the compensation is also evident in the functional connectivity networks of specific brain regions (Fig. 4), indicating a more complex relationship between neuronal and neuromodulatory activities beyond simple correlation^15^. By analyzing the low-dimensional manifolds of odor identity representation, we found ACh is not only involved in odor representation, but also show more stable representation across several hours than neuronal activity (Figs. 3-5). While recent studies have considered the spatiotemporal changes in odor representation^23,26,53^, our findings provide the first extensive description of olfactory representation across such a large temporal and spatial scale, encompassing both neuronal and neuromodulatory activities, paving the way for a more comprehensive understanding of neural information representation.

Traditionally, information is considered coded in neuronal activity and can be decoded with high precision^64^. Neuromodulators and neurotransmitters have been primarily understood as neuronal activity regulators rather than direct information carriers. However, the recent evidence of dissociable dopamine firing and release and sensory-related fast cholinergic activities prompts a reevaluation of neuronal and neuromodulatory information representation^15,17,28,65^. Our study has directly uncovered the compensation of olfactory information representation between neuronal activity and ACh dynamics, as well as the superior long-term representational stability of ACh. These findings highlight the prominent involvement of ACh in shaping olfactory representation and emphasize the need for more than just considering neuronal activity when examining information representation of the brain. Such involvement can be further investigated in more complex tasks in the future.

Since we used pan-neuronally labeled flies to perform a screen in this study, we could not resolve single neurons or distinguish neuron types. Further investigation of genetically targeted circuits and neuron types is necessary to elucidate the mechanisms of the local mismatch between the neuronal and neuromodulatory activities as well as the global compensation. Large-scale sparse labeling can also greatly assist in this endeavor^49,66^. Additionally, the extensive-surveying methodology employed in our study may hold promise for uncovering new findings in other areas, such as learning and memory, which have widespread effects across the brain. Moreover, the simultaneous volumetric recording of multiple indicators on a large scale also presents valuable opportunities to investigate the interactions between different neuromodulators^67,68^ and the coupling of other important processes^69^.

## Methods

### Fly stocks

Flies with pan-neuronal expression of jGCaMP7f and rGRAB_ACh-0.5 were of the genotype: w; UAS-rGRAB_ACh-0.5/+; nSyb-Gal4, UAS-jGCaMP7f, which was combined by w; UAS-rGRAB_ACh-0.5/cyo; +/(TM2/TM6B), nSyb-Gal4 and UAS-jGCaMP7f. Flies with pan-neuronal expression of jGCaMP7f and rGRAB_HTR2C-0.5 were of the genotype: w; UAS-rGRAB_HTR2C-0.5/+; nSyb-Gal4, UAS-jGCaMP7f, which was combined by w; UAS-rGRAB_HTR2C-0.5/cyo; +/(TM2/TM6B), nSyb-Gal4 and UAS-jGCaMP7f. w; UAS-rGRAB_ACh-0.5/cyo; +/(TM2/TM6B) and w; UAS-rGRAB_HTR2C-0.5/cyo; +/(TM2/TM6B) were from Yulong Li’s Lab at Peking University. nSyb-Gal4 (BDSC: 51941) and UAS-jGCaMP7f^32^ were from Yi Zhong’s Lab at Tsinghua University.

### Odor delivery

3-octanol, 4-methylcyclohexanol, and ethyl acetate diluted 1.5:1000, 1:1000, 1:1000 in mineral oil were used as odors^70^. Odors were delivered for 5 s with 30 s inter-stimuli intervals for 180 trials (60 sessions) using a custom-made air pump, with a pseudo-random order, and avoiding consecutive presentations of the same odor.

### Fly preparation for functional imaging

Flies were raised on standard cornmeal medium with a 12-h light/12-h dark cycle at 23 °C and 60% humidity and housed in mixed male/female vials. 3-8-day-old female flies were selected for brain imaging. To prepare for imaging, flies were anesthetized on ice and mounted in a 3D-printed plastic disk that allowed free movement of the legs, as previously reported^70,71^. The posterior head cuticle was opened using sharp forceps (5SF, Dumont) at room temperature in fresh saline (103 mM NaCl, 3 mM KCl, 5 mM TES, 1.5 mM CaCl_2_, 4 mM MgCl_2_, 26 mM NaHCO_3_, 1 mM NaH_2_PO_4_, 8 mM trehalose, and 10 mM glucose (pH 7.2), bubbled with 95% O_2_ and 5% CO_2_)^70,71^. After that, the fat body and air sac were also removed carefully. The position and angle of the flies were adjusted to keep the posterior of the head horizontal, and the window was made big and clean, for the convenience of multiple-brain-region observation. Brain movement was minimized by adding UV glue around the proboscis^51,72^. After preparation, flies were placed under the objective for two-photon imaging.

### Multiple-brain-region two-photon volumetric imaging by 2pSAM

We used a 25×/1.05 NA water immersion objective (Olympus) in this experiment. Min-NA 2pSAM was adopted to achieve an effective depth of focus of 100 μm^31^, covering about half of the axial range of the fly brain (∼ 80 - 180 μm under the surface). The field of view was 458.7 μm × 458.7 μm (512 pixels × 512 pixels), covering the whole lateral range of the central brain and some part of the optic lobe. An excitation wavelength of 1035 nm was used for two-color imaging. The power of the excitation light was set at 25 - 35 mW through the ∼ 2 h recording. For detection, a 525-nm filter (MF525-39, Thorlabs) was used for the green channel, and a 610-nm filter (ET610/75m, Chroma) was used for the red channel. The acquiring rate was 30 Hz, and a 13-angle scanning was adopted.

### Image processing pipeline

#### Preprocessing - registration and denoising

We conducted preprocessing steps for the images of each angle. First, we performed motion correction using a piecewise rigid registration algorithm as previously described^31,73^. Then, we applied denoising algorithms to enhance the signal-to-noise ratio, which was essential for our voxel-based analyses. We utilized DeepCAD-RT in most of our analyses due to its superior capability in detecting subtle signal changes^35,36^. To improve the recovery of temporal characteristics in manifold analyses, we implemented a new algorithm SRDTrans^37^. We trained customized denoising models for each channel of each fly.

#### Volume reconstruction

We utilized VCD-Net for reconstruction to accelerate the process^74^. First, we sampled about 2% of the frames for reconstruction using the previously reported algorithm^31^. The reconstructed volumes paired with the light-filed images served as training sets. Subsequently, all the volumes of an experiment were efficiently reconstructed using VCD-Net within a few hours. The reconstructed volumes were 256 pixels × 256 pixels × 25 pixels, with a lateral resolution of 1.79 μm and an axial resolution of 4 μm. This resolution was decided to strike a balance between achieving high resolution and managing the total data size for subsequent analyses. The temporal rate of volumes was 30 Hz and 2.3 Hz with and without the sliding-window reconstruction method, respectively.

#### Alignment to atlas

The *Drosophila* brain atlas was sourced from Virtual Fly Brain, the version used in a previous work^42^. The alignment was performed using landmarks with ImageJ (https://imagej.net/plugins/name-landmarks-and-register)^51^. We used the red channel for registration, as the structure of this channel was more distinct. An eroded atlas, in which the mask of each region was eroded, was used to extract brain regions, preventing incorrect assignment of edge voxels to regions. While completing volume registration in the previous steps, we only aligned the first volume to the atlas, thus aligning the entire stack.

#### Temporal trace extraction

We extracted the temporal traces of each voxel. For the computation of Δ*F/F*, a sliding window was used. For each frame, the mean of the lower 30% intensity of the previous 200 frames (at 2.3Hz) was taken as *F_0_*, and Δ*F/F* was computed as (*F-F_0_*)*/F_0_*, in which *F* was the current intensity of the voxel. The Δ*F/F* of each brain region was taken as the average of the Δ*F/F* of the voxels within the region. Since the odor responses exhibited persistence beyond the duration of odor delivery, we extended the time window for extracting these responses. Specifically, the selected time windows encompassed the period from the onset of the response to its decay to less than half of the peak magnitude. In most of our analyses, as the dynamics of r5-HT were slower, the time windows were approximately 8 s, 8 s, and 16 s (20 frames at 2.3 Hz, 2.3 Hz, and 1.15 Hz) for G7f, rACh, and r5-HT, respectively. In the case of odor identity classification and manifold analyses, the time windows were set to approximately 12 s (15 frames at 1.15 Hz) and 33 s (40 frames at 1.15 Hz) after odor delivery for all indicators.

### The map of responsiveness

For each fly, the Pearson correlation between trial-averaged Δ*F/F* and stimulus (binary sequence) was calculated for each voxel in each channel to obtain the maps of responsiveness (Fig. 2a). As there was a delay between stimulus and response, we shifted the stimulus window forward and took the maximum of the correlation as the result value. The maps of individual flies were aligned and averaged to generate a summary map of the sample flies. The correlation of each region was the average of the voxels within the region (Extended Data Fig. 1b).

### The map of response intensity

For each trial, the standard deviation of Δ*F/F* in 8 s (G7f and rACh) and 16 s (r5-HT) since odor delivery was calculated as the response intensity. To create a map, we calculated the average response intensity for each voxel across trials. The maps of individual flies were aligned and averaged to generate a summary map of the sample flies, as shown in Fig. 2a. The response intensity of each region was the average of the voxels within the region (Extended Data Fig. 1c).

### Phase delay and pulse width

For each fly, the phase delay and pulse width were calculated for each voxel in each channel using the trial-averaged Δ*F/F* (Fig. 2d). The phase delay was the time-lapse from the start to the peak of the response (Extended Data Fig. 2a). The pulse width was the full width at half maximum (Extended Data Fig. 2a). The maps of individual flies were aligned and averaged to generate a summary map of the sample flies. The phase delay and pulse width of each region were the average of the voxels within the region (Fig. 2g, h; and Extended Data Fig. 2b, e). To visualize the distribution of phase delay (or pulse width) of rACh or r5-HT relative to G7f of the voxels in each region, we consolidated the data of all flies and plotted the histogram of the phase delay (or pulse width) difference (Extended Data Fig. 2c, d, f, and g). By sorting the phase delay of each brain region, we could yield a schematic of the global information propagation for each indicator (Fig. 2i).

### Brain-region-level functional connectivity network analysis

Brain-region-level functional connectivity networks of the left-side brain regions and the central complex recorded (23 brain regions) were analyzed.

#### Construction of functional connectivity networks

To construct a brain-region-level functional connectivity network, we first spliced the response records for each voxel over the time window of 180 trials. We then averaged all voxels within brain regions to obtain the average response record for each brain region and computed the Pearson correlation of the averaged response records between each pair of brain regions. The above process was performed for each indicator, such that the correlation matrices shaped in 23 × 23 dimensions were generated for G7f, rACh, and r5-HT, respectively. Each correlation matrix was used to construct a weighted undirected network (Fig. 2j; and Extended Data Fig. 3a, b), where the nodes represented brain regions, and the edges represented the correlation values between brain regions. We used the Louvain and greedy algorithms to detect the community structure in the functional connectivity networks and obtained consistent results. The detected communities formed by the tightly connected brain regions were labeled with dashed circles (Fig. 2j and Extended Data Fig. 3a).

#### Comparison of functional connectivity patterns in physical space

We examined the changes in the distribution of functional connectivity in physical space using the nonparametric multifactor ANOVA test called the Scheirer–Ray–Hare test. Specifically, we arranged the functional connections of each fly during odor stimulation and the resting state in the same order and grouped the data from multiple flies, thus testing whether odor stimulation was a factor that significantly affected the brain-region-level functional connectivity (Fig. 2k). In the same way, sorting and grouping data from G7f, rACh, and r5-HT allowed testing whether the indicators significantly affected functional connectivity. Fig. 2k displayed the functional connectivity between each pair of brain regions averaged across multiple flies, sorted by the connection strength of G7f in the resting state.

#### Comparison of functional connectivity in network topology

We analyzed the distinctions in the topological characteristics of the brain-region-level functional connectivity networks across different indicators and states. The degree of a node was the total connection strength of the edges connected, and the average degree of a network represented the average degree across all nodes. To assess the differences in the distribution of node degrees (Fig. 2l and Extended Data Fig. 3e) and the average degree values (Extended Data Fig. 3c, f) across different indicators and states, we used the Kolmogorov-Smirnov test and the Wilcoxon signed-rank test, respectively. Additionally, we calculated the “community ratio”, which was defined as the connection strength ratio of the community containing most olfactory regions, to reflect the significance of the community (Fig. 2m and Extended Data Fig. 3d). A larger community ratio indicated a more influential role of the given community in the network.

### Accuracy map

The volume was divided into small blocks of 4 × 4 × 2 voxels. The data form within each block was trials × [voxels × frames × channels]. We first performed PCA and kept the dimensions explaining 90% variance. Then, SVM was applied to classify the trials according to odor identities, and the accuracy was obtained by 5-fold cross-validation. The maps of individual flies were aligned and averaged to generate a summary map of the sample flies (Fig. 3a and Extended Data Fig. 1d). The accuracy of each region was the average of the voxels within the region (Fig. 3b and Extended Data Fig. 1d). The map revealing the accuracy improvement was generated by subtracting the accuracy map of G7f from the accuracy map of the dual-channel data (Fig. 4a and Extended Data Fig. 8a). We employed PCA and SVM on frames 1-14 from the start of odor delivery.

### Voxel-level odor identity classification

#### Odor identity classification in each brain region

The data within each region was organized in a [trials × frames] × [voxels × channels] format. To reduce the dimensionality, we initially used PCA. Next, we employed LDA to identify a stable low-dimensional space where odor identity was well-represented. The data was then reshaped as trials × [frames × LDA dimensions]. SVM was utilized for trial classification, and the accuracy was determined by 5-fold cross-validation. It was worth noting that the classification accuracy varied based on the variance threshold used in the PCA step (Extended Data Fig. 4a, b). A threshold of 0.8 exhibited optimal performance and was selected. We employed PCA and SVM on frames 1-14, and LDA on frame 3 from the start of odor delivery.

#### Odor identity classification across multiple brain regions

First, for single-channel classification, a brain-region mask was applied to selectively choose the voxels within the community containing olfactory regions. For dual-channel classification, the brain-region mask for the rACh or r5-HT channel was the brain regions with accuracy gain, either in the average accuracy or the accuracy of the voxel-level classification (Fig. 4b; and Extended Data Figs. 5d and 8b). Then, to handle the higher dimensionality of multiple-brain-region data compared to single-region data, we implemented an additional block-wise PCA step prior to the steps of *odor identity classification in each brain region*. The volume was divided into 10 × 10 × 10 blocks. Within each block, PCA was carried out while retaining dimensions accounting for 90% of the variance. When dealing with dual-channel data, this approach was independently applied to each channel, and the resulting outputs were consolidated for the subsequent steps. We observed that the classification accuracy varied with the number of dimensions retained after the second PCA step (Extended Data Fig. 4c-f). Interestingly, the accuracy became stable after surpassing a certain threshold of dimensions. Hence, we set the threshold at 25 for optimal results. We employed PCA and SVM on frames 1-14, and LDA on frame 3 from the start of odor delivery. We utilized the two denoising algorithms, SRDTrans (Extended Data Fig. 4c, d) and DeepCAD-RT (Extended Data Fig. 4e, f), and obtained similar results and robust accuracy gain. The results of SRDTrans were analyzed.

### Brain-region-level odor identity classification

The average Δ*F/F* of single brain regions were consolidated for classification. The steps were similar to *odor identity classification in each brain region*. The variance threshold of the PCA step was set to 0.998 for optimal performance (Extended Data Fig. 4g, h).

### Manifolds

For the odor identity classification across multiple brain regions, we employed PCA and SVM on frames 1-14, and LDA on frame 3 from the start of odor delivery. Furthermore, we applied the PCA and LDA transformations to all frames to capture the low-dimensional dynamics in the LDA space. This approach allowed us to observe that the responses of individual trials within this low-dimensional space manifested as curved traces, which collectively formed a manifold. We obtained the manifold from 2 s before the onset of odor stimuli to 33 s thereafter to characterize the whole process of odor responses, starting from and returning to a random state. Each trace started from the common state and extended in different directions, symbolizing the essence of various odor identities. To visualize the process of extending and returning clearly, we averaged the traces with the same odor identity and colored the average trace with the time-lapse relative to odor delivery (Figs. 3d-f, 4d, e; and Extended Data Fig. 8d, e).

### Manifold analysis

We aligned and combined the manifolds of the testing sets in each fold and examined the characteristics of the combined manifold. The average traces of each odor identity were used to measure the distance to the origin and the inter-class distance. The distance to the origin referred to the distance between the locations of a specific time point and the average location of the start time point (2 s before the onset of odor stimuli) in the LDA space. The inter-class distance referred to the distance of odor pairs (O-M: OCT and MCH, O-E: OCT and EA, M-E: MCH and EA) at each time point. The intra-class distance referred to the average distance between the traces with the same odor identity at each time point. The return time was evaluated as the time-lapse for distance to the origin or intra-class distance to recover to the random level (the average value of 29-30 s) from the start of odor stimuli. The average intra-class distance was calculated within 12 s after odor delivery. The values of each odor were compared to analyze the representational distinctions among odor identities (Extended Data Fig. 7). The average value of all odors was evaluated to analyze the representational distinctions among indicators (Figs. 3 and 4; and Extended Data Fig. 8). A fly co-labeled by G7f and rACh and a fly co-labeled by G7f and r5-HT are excluded in this analysis for the alignment failure of each fold.

### Functional connectivity network analysis within brain regions

#### Construction of functional connectivity networks within brain regions

To construct a functional connectivity network within each brain region, we first spliced the response records for each voxel of a given brain region over the time window of 180 trials. We then computed the Pearson correlation of the response records between each pair of voxels and generated an N × N correlation matrix, N referring to the number of voxels in a given brain region. Thus, a G7f matrix and a rACh matrix were obtained for each brain region of each fly co-labeled by G7f and rACh (Fig. 4i, j); a G7f matrix and a r5-HT matrix were obtained for each brain region of each fly co-labeled by G7f and r5-HT (Extended Data Fig. 8i, j). We used all the N voxels of a given brain region as nodes and retained the top 30% correlations in the matrix as edges, thus obtaining a weighted undirected network for each region. We displayed the constructed network of each brain region according to the relative position of each voxel in physical space, where the edges were shown in gray, and the nodes were colored by the community division (using the Louvain algorithm) of the network (Fig. 4i, j; Extended Data Figs. 8i, j, and 9a, b). Seven brain regions were selected from different neuropils of the fly brain (Supplementary Table 1) to demonstrate their functional correlation matrices and networks during odor stimulation (Extended Data Fig. 9a) and the resting state (Extended Data Fig. 9b) for rACh and G7f.

#### Comparison of functional connectivity in physical space

In order to compare the neuromodulatory (rACh or r5-HT) and neuronal (G7f) functional connectivity, we calculated the difference matrix Δ as follows:

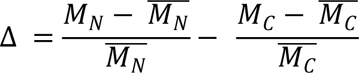

Where *M_C_* and 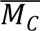 denote the correlation matrix of G7f and its average value, *M_N_* and 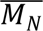 denote the correlation matrix of rACh or r5-HT and its average value (Fig. 4i, j; Extended Data Figs. 8i, j, and 9a, b). The neuromodulatory and difference matrices were sorted according to the clustering of the G7f matrix, so that the black boxes marked the most prominent functional connections in the G7f matrix, while the outside of the black boxes demonstrated the correlations between different clusters (Fig. 4i, j). The red boxes in Fig. 4i signed the functional connections highlighted by the rACh matrix that were not emphasized in the G7f matrix, suggesting the connectivity compensation. We showed the nodes involved in around top 1% correlations of the difference matrix Δ in coral color and the remaining nodes in blue in the networks (Fig. 4i, j; Extended Data Figs. 8i, j, and 9a, b). This reflected whether the connections emphasized by neuromodulators gathered together in physical space. We designed a metric 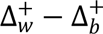 to measure connectivity compensation. The metric 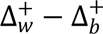 was defined as the difference between the mean of positive values within and between clusters in the difference matrix Δ. The results showed that this metric was always greater than 0. The smaller the metric 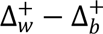, the more relative connectivity compensation there was (Fig. 4k and Extended Data Fig. 8k). All nine brain regions with accuracy gain and four without accuracy gain were compared for rACh. The Wilcoxon signed-rank test was used to detect whether the metric 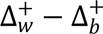 of each brain region with accuracy gain was significantly less than the average level of the selected brain regions without accuracy gain, i.e., LH, EPA, AOTU, and IVLP. Although for r5-HT, only MBVL exhibited accuracy gain (Extended Data Fig. 5d), we performed the same test as rACh.

#### Comparison of functional connectivity in network topology

In order to compare the topological characteristics between neuronal and neuromodulatory functional connectivity networks, we measured the average clustering coefficient and the node centrality (degree centrality) distribution. The clustering coefficient was a measure of the degree to which nodes clustered together, and the average clustering coefficient of the network was the average of the local clustering coefficients of all nodes, i.e., 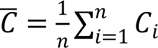 referring to the number of nodes. We calculated the average clustering coefficients of the functional connectivity networks for different indicators in brain regions with and without accuracy gain (Fig. 4l and Extended Data Fig. 8l). The larger the average clustering coefficient was, the stronger the nodes in the functional connectivity network clustered together locally. We then computed the difference Δ*_Clustering_* between the average clustering coefficients of neuronal and neuromodulatory functional connectivity networks (Fig. 4m and Extended Data Fig. 8m). We also calculated the probability density function (PDF) and cumulative distribution function (CDF) of the degree centrality of nodes in neuronal and neuromodulatory functional connectivity networks in brain regions with and without accuracy gain (Fig. 4n; Extended Data Figs. 8n and 9c). A single-peaked PDF indicated that most nodes in the network had similar degree centrality, in contrast, a bimodal distribution indicated that the nodes in the network had more heterogeneity in their attributes. The difference between the degree centrality distributions of different indicators was measured by the Wasserstein distance^75^ (Fig. 4o and Extended Data Fig. 8). We also used another distance function called the Energy distance and obtained similar results^76^. The Wilcoxon signed-rank test was used to detect whether the above topology properties of each brain region with accuracy gain for rACh were significantly higher or lower than the average level of the selected brain regions without accuracy gain, i.e., LH, EPA, AOTU, and IVLP. Although for r5-HT, only MBVL exhibited accuracy gain (Extended Data Fig. 5d), we performed the same test as rACh.

### Temporal changes of manifolds

To analyze the temporal changes, the 60 sessions throughout the experiment were divided evenly into four stages, i.e., S1: Sessions 1-15, S2: Sessions 16-30, S3: Sessions 31-45 and S4: Sessions 46-60. We aligned and combined the manifolds of each fly to facilitate the following analyses. The low-dimensional traces of the corresponding sessions formed the manifolds of each stage. The characteristics of the manifolds at each stage were assessed. Distance to the origin, inter-class distance, and intra-class distance were measured as described above. Additionally, we examined the return locations (the average location of 29-30 s from the start of odor stimuli) of the sessions. We measured the x- and y-coordinates of the return locations and the distances between the return locations and the average location of the start time point (2 s before the onset of odor stimuli). Intra-class distance was measured for every three consecutive sessions, and other metrics were measured for every session. The average values of each stage were compared (Fig. 5a-e; and Extended Data Fig. 10a, b).

### Temporal changes of functional connectivity networks

The brain-region-level functional connectivity networks for each stage were generated using the abovementioned method. We examined changes in the distribution of functional connectivity in physical space over time using the Scheirer–Ray–Hare test (Fig. 5g), and evaluated changes in the topological features of the functional connectivity networks. The changes in degree distribution (Fig. 5h and Extended Data Fig. 10c) and average degree values (Fig. 5i) of the functional connectivity networks were examined by the Kolmogorov-Smirnov test and the Wilcoxon signed-rank test, respectively. We further examined whether the community structure of the functional connectivity networks for each indicator changed significantly over time by calculating the community ratio (Fig. 5j).

### Motion analysis

We captured videos of fly abdomens and extracted the motions by computing the absolute differences between consecutive frames (Extended Data Fig. 6a). The resulting motion energy, averaged across the FOV, was then correlated with the stimulus presented to the flies (binary sequence) using Pearson correlation (Extended Data Fig. 6b). To match the temporal resolution and periods of the neuronal and neuromodulatory activities, we downsampled the videos and extracted 12 s time windows from the 180 trials. PCA was applied to reduce the dimensionality and extract the behavioral features (Extended Data Fig. 6c). We retained the first 50 PCs. SVM was utilized to classify stimulus periods and intervals as well as odor identities based on these behavioral features (Extended Data Fig. 6d). Behavior during odor stimulation (5 s) was used in the classification. We conducted a label shuffling as a negative control and evaluated accuracies using 5-fold cross-validation. Next, we used Ridge regression to predict the first 30 PCs of the multiple-brain-region neuronal and neuromodulatory activities from behavior, stimulus, or both (Extended Data Fig. 6e-m). “Stimulus” refers to the trial-averaged responses to each odor identity. R^2^ was assessed by 5-fold cross-validation. We used the partial PCs explained by behavior and the residual after subtracting this explained part to conduct odor identity classifications (Extended Data Fig. 6g, j, and m). The first 30 PCs were employed, following the same classification method as using the behavioral features.

## Acknowledgments

We thank Jun Zhou, Han Mo, Yunchuan Zhang, Wantong Hu, and Qi Yang at Yi Zhong’s lab, and Xuelin Li at Yulong Li’s lab for their help with *Drosophila* experiments, and Yeyi Cai, Yuanlong Zhang, and Tao Sun at our lab for their help with data processing. The figures of *Drosophila* are created with BioRender.com.

## Author contributions

Conceptualization: JW, LF, QD, JF; Methodology: JF, JW, YW, LL, ZZ, JH, GL, FD, XL, YL; Investigation: JF, YW, LL, ZZ, YZ, JZ, XH; Visualization: JF, YW, LL; Funding acquisition: QD, LF, JW; Project administration: QD, LF, JW; Supervision: QD, LF, JW; Writing – original draft: JF, JW, LL; Writing – review & editing: QD, LF, YL, YW.

## Data and code availability

The demo data of an example fly is available on <u>OneDrive</u>. The codes for data analysis are available on <u>GitHub</u>. We will upload the final version of the data and codes on the data platforms like Zenodo after revision. The entire dataset with a total size of 5 TB, which includes the extracted neuronal and neuromodulatory traces within the 3D volumes over 2h of 10 flies co-labeled by G7f and rACh and 10 flies co-labeled by G7f and r5-HT, will be open-sourced after publication, as an important resource for the neurobiology and computational neuroscience communities.

---

**Extended Data Fig. 1.**
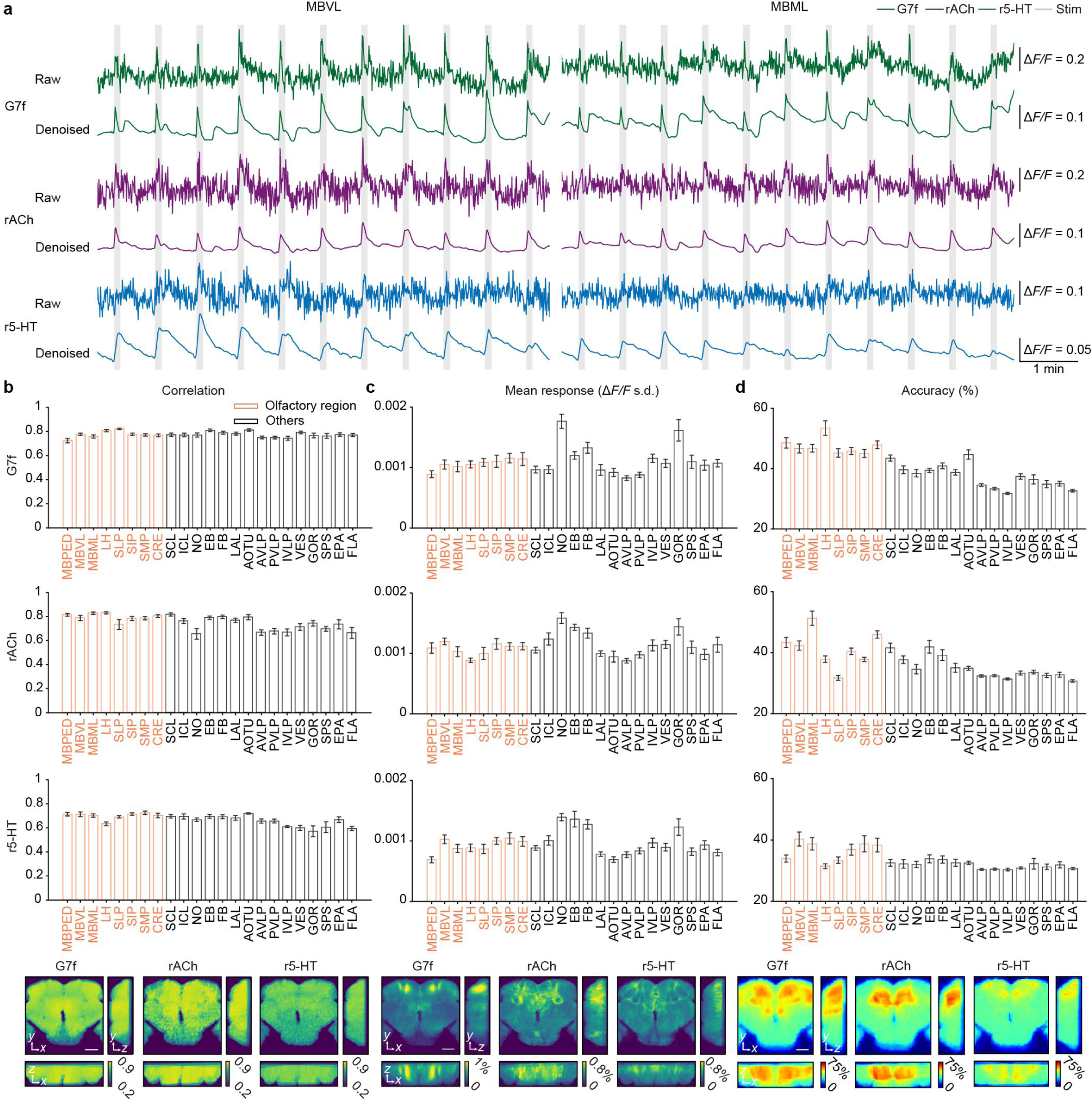
The performance of denoising and the statistics of responsiveness, response intensity, and odor identity classification accuracy in each brain region. **a**, The average Δ*F/F* of two brain regions for three indicators of an example fly, before and after the denoising by DeepCAD-RT. **b**-**d**, Top: The average correlation between trial-averaged G7f, rACh, or r5-HT activities and odor stimulus (**b**, responsiveness), the average standard deviation of Δ*F/F* during odor stimulation across trials (**c**, response intensity), and the average odor identity classification accuracy (**d**) in each region. Bottom: The corresponding maps averaged across flies. Based on the statistical data, it is evident that while the level of responsiveness or response intensity in olfactory regions is not notably superior to that of non-olfactory regions, the odor identity classification accuracy of most olfactory regions surpasses other regions obviously, especially for G7f. For rACh and r5-HT, there are olfactory regions with low responses as well as low accuracies (LH, SLP, MBPED), in contrast to G7f. 10 flies co-labeled by G7f and rACh and 10 flies co-labeled by G7f and r5-HT are analyzed. *n* = 20 flies for G7f, *n* = 10 flies for rACh, *n* = 10 flies for r5-HT, mean ± s.e.m. A fly co-labeled by G7f and rACh with an obvious lean angle of the brain is excluded for clarity in **b**. Flies co-labeled by G7f and rACh with high-intensity non-specific fluorescence on the upper edge of the brain are excluded for clarity in **c**. Results of the left-side brain regions and the central complex are shown in the statistical charts. Scale bars: 50 μm in **b**-**d**.

**Extended Data Fig. 2.**
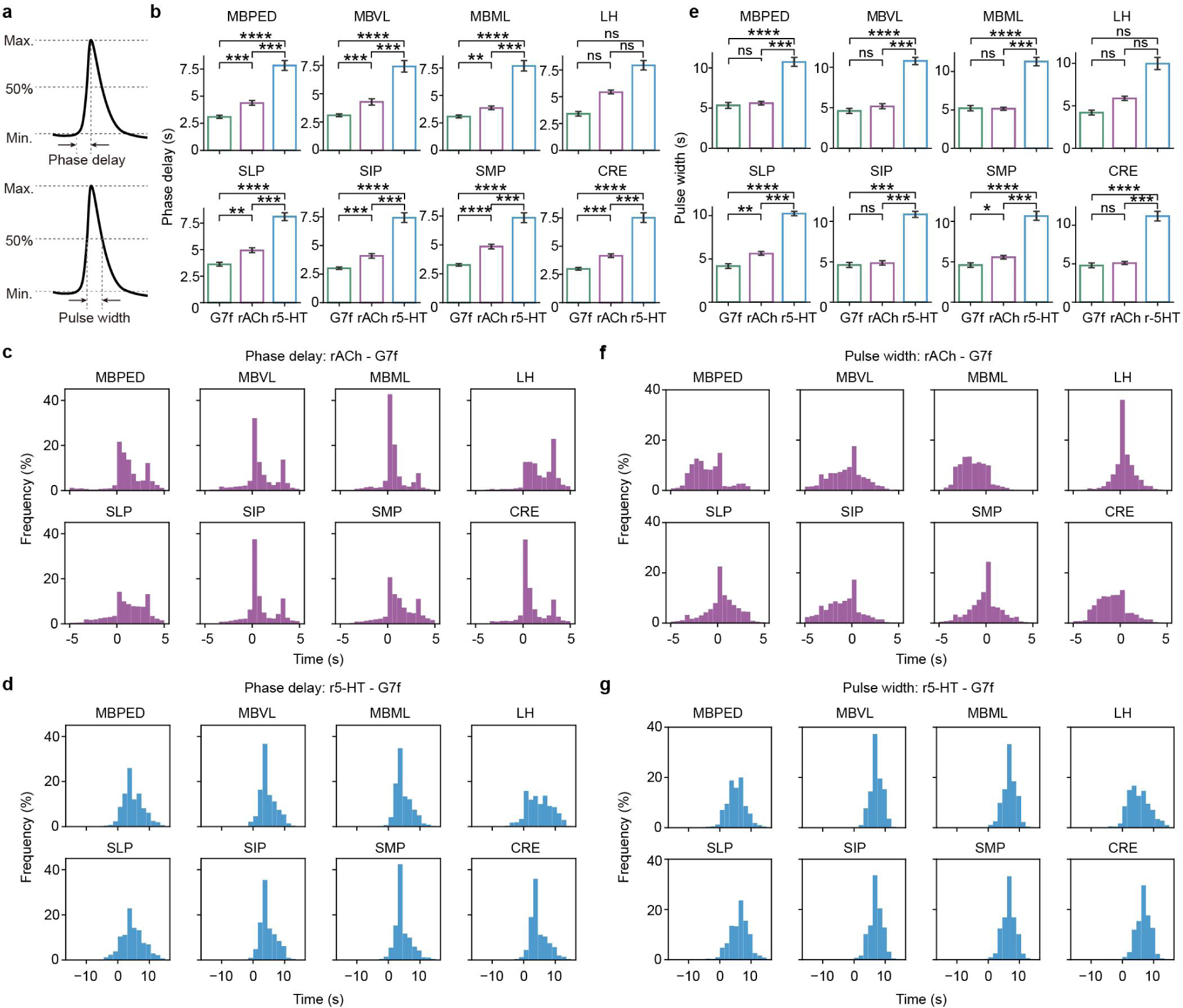
Phase delay and pulse width of the odor responses in olfactory regions. **a**, The schematic of the calculation of phase delay and pulse width. **b**, Statistics of the average phase delay of the odor responses for three indicators in each olfactory region. **c**, Distribution of the phase delay difference between rACh and G7f of the voxels in each region, consolidating the data of all flies. **d**, Similar to **c**, but for r5-HT and G7f. **e-g**, Similar to **b**-**d**, but for pulse width. 10 flies co-labeled by G7f and rACh and 10 flies co-labeled by G7f and r5-HT are analyzed. *n* = 20 flies for G7f, *n* = 10 flies for rACh, *n* = 10 flies for r5-HT, mean ± s.e.m in **b**, **e**. *n* = 10 flies in **c**, **d**, **f**, and **g**. A fly co-labeled by G7f and rACh with the interference of the calculation of phase delay by the non-specific fluorescence on the upper edge of the brain is excluded in **b**, **c**, **e**, and **f**. Flies without a general coverage of region LH are excluded from the statistics of this region. Results of each olfactory brain region in the left semi-brain are shown. Two-sided Mann-Whitney U test in **b**, **e**, \*\*\*\**P* < 0.0001, \*\*\**P* < 0.001, \*\**P* < 0.01, \**P* < 0.05, ns - not significant (*P* > 0.05).

**Extended Data Fig. 3.**
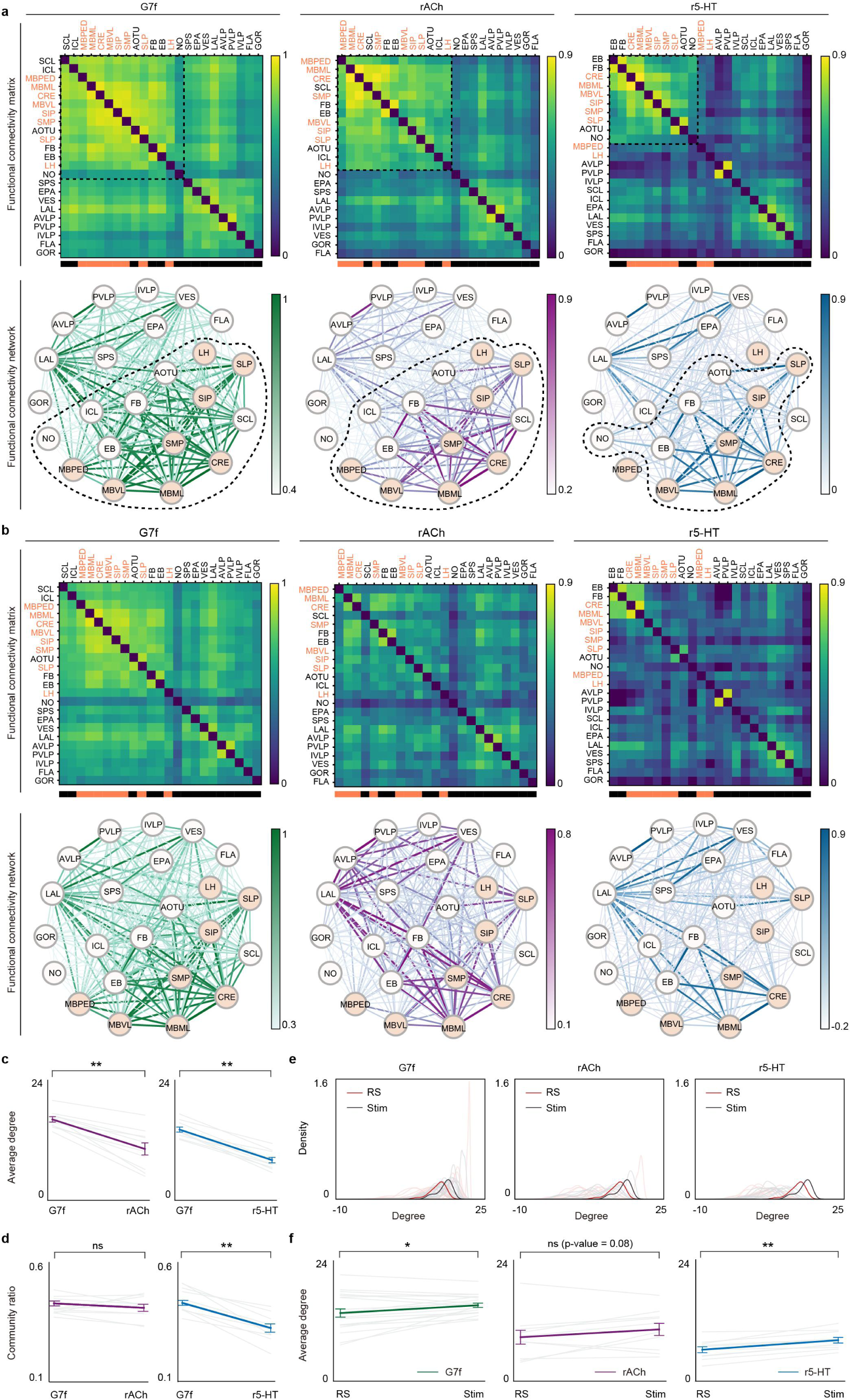
The functional connectivity networks of three indicators during different states show distinct properties. **a**, Clustered functional connectivity matrices and networks of the left-side brain regions and the central complex for each indicator during odor stimulation, consistent with Fig. 2j. The names of the brain regions are represented here. **b**, Functional connectivity matrices and networks of the left-side brain regions and the central complex for each indicator in the resting state. The order of the matrix elements and the placement of the network nodes are consistent with **a**. **c**, **d**, The average degree and community ratio of the functional connectivity networks for different co-labeled indicators. **e**, The PDF of the average degree in the functional connectivity networks for three indicators during odor stimulation (black lines) and the resting state (red lines). **f**, The statistics of the average degree of the functional connectivity networks during different states. In **c**-**f**, each light-colored line represents a fly.10 flies co-labeled by G7f and rACh and 10 flies co-labeled by G7f and r5-HT are analyzed. *n* = 20 flies for G7f, *n* = 10 flies for rACh, *n* = 10 flies for r5-HT, mean ± s.e.m in **a**, **b**, **e**, and **f**. *n* = 10 flies, mean ± s.e.m in **c**, **d**. Stim: Odor stimulation. RS: The resting state. Two-sided Wilcoxon signed-rank test; \*\*\*\**P* < 0.0001, \*\*\**P* < 0.001, \*\**P* < 0.01, \**P* < 0.05, ns - not significant (*P* > 0.05).

**Extended Data Fig. 4.**
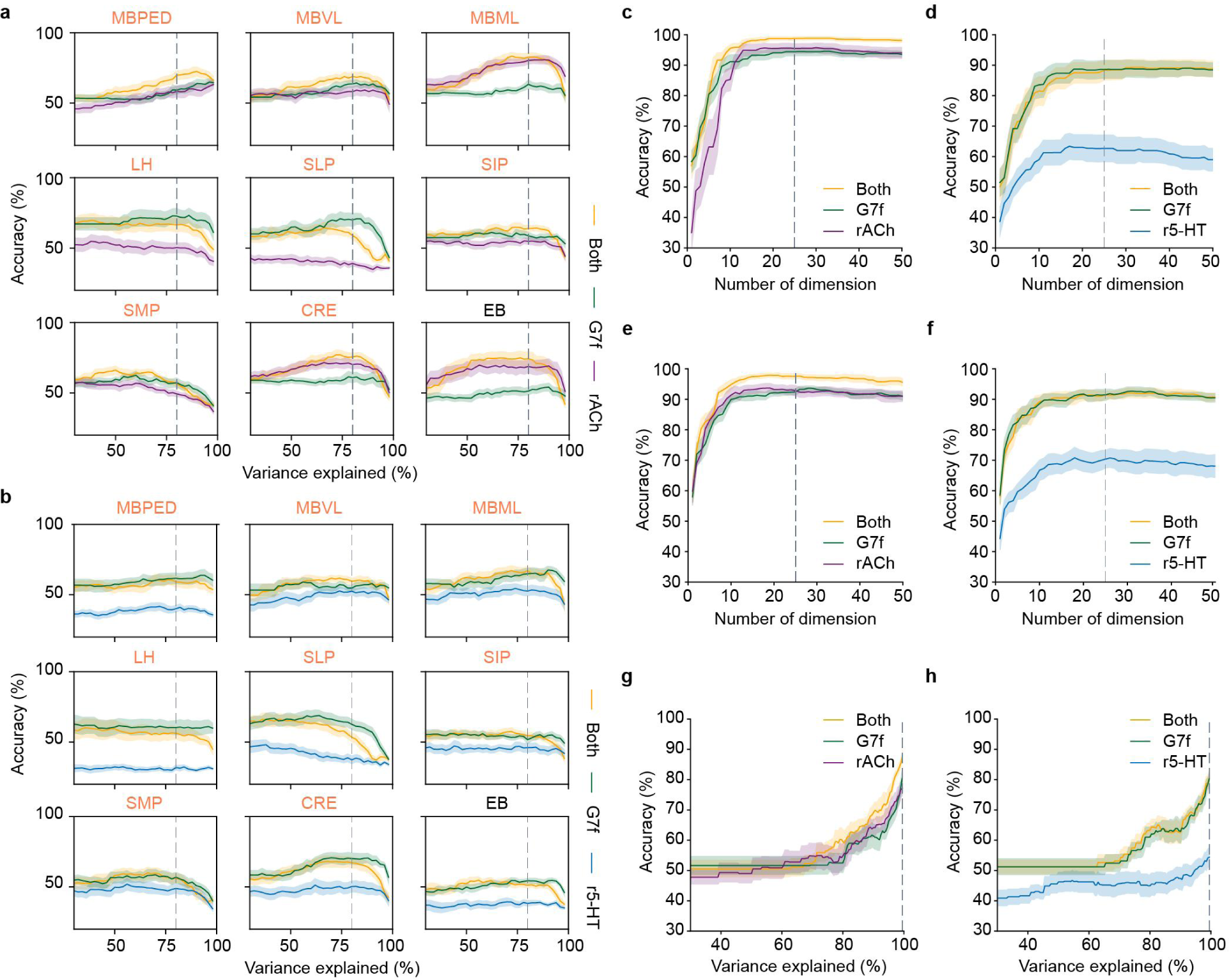
PCA threshold determination for odor identity classifications. **a**, The accuracy of the voxel-level odor identity classification in 9 brain regions (left-side olfactory brain regions and EB) changing with the explained variance of the PCs retained, for G7f, rACh, and integrating both channels. The accuracy does not show significant change with the explained variance. A threshold of 0.8 is taken as most brain regions reach a high and stable level of accuracy, marked by the dashed line. **b**, Similar to **a**, but for flies co-labeled by G7f and r5-HT. **c**, The accuracy of the voxel-level multiple-brain-region odor identity classification changing with the dimensions of PCs retained (Step 3 in Fig. 3c), for G7f, rACh, and integrating both channels. Integrating both channels yields a higher accuracy for almost all dimension values. The dashed line (dimension = 25) marks the threshold taken, as the average accuracy reaches a high and stable level for all indicators. **d**, Similar to **c**, but for flies co-labeled by G7f and r5-HT. **c**, **d**, are results using the denoising algorithm SRDTrans. **e**, **f**, Similar to **c**, **d**, but are results using the denoising algorithm DeepCAD-RT. Obvious accuracy gains for integrating rACh and G7f signals exist using both denoising algorithms. **g**, The accuracy of the region-level multiple-brain-region odor identity classification changing with the explained variance of the PCs retained, for G7f, rACh, and integrating both channels. A threshold of 0.995 is taken, marked by the dashed line. **h**, Similar to **g**, but for flies co-labeled by G7f and r5-HT. 10 flies co-labeled by G7f and rACh and 10 flies co-labeled by G7f and r5-HT are analyzed. *n* = 10, mean ± s.e.m (shades). Flies without a general coverage of region LH are excluded from the statistics of this region.

**Extended Data Fig. 5.**
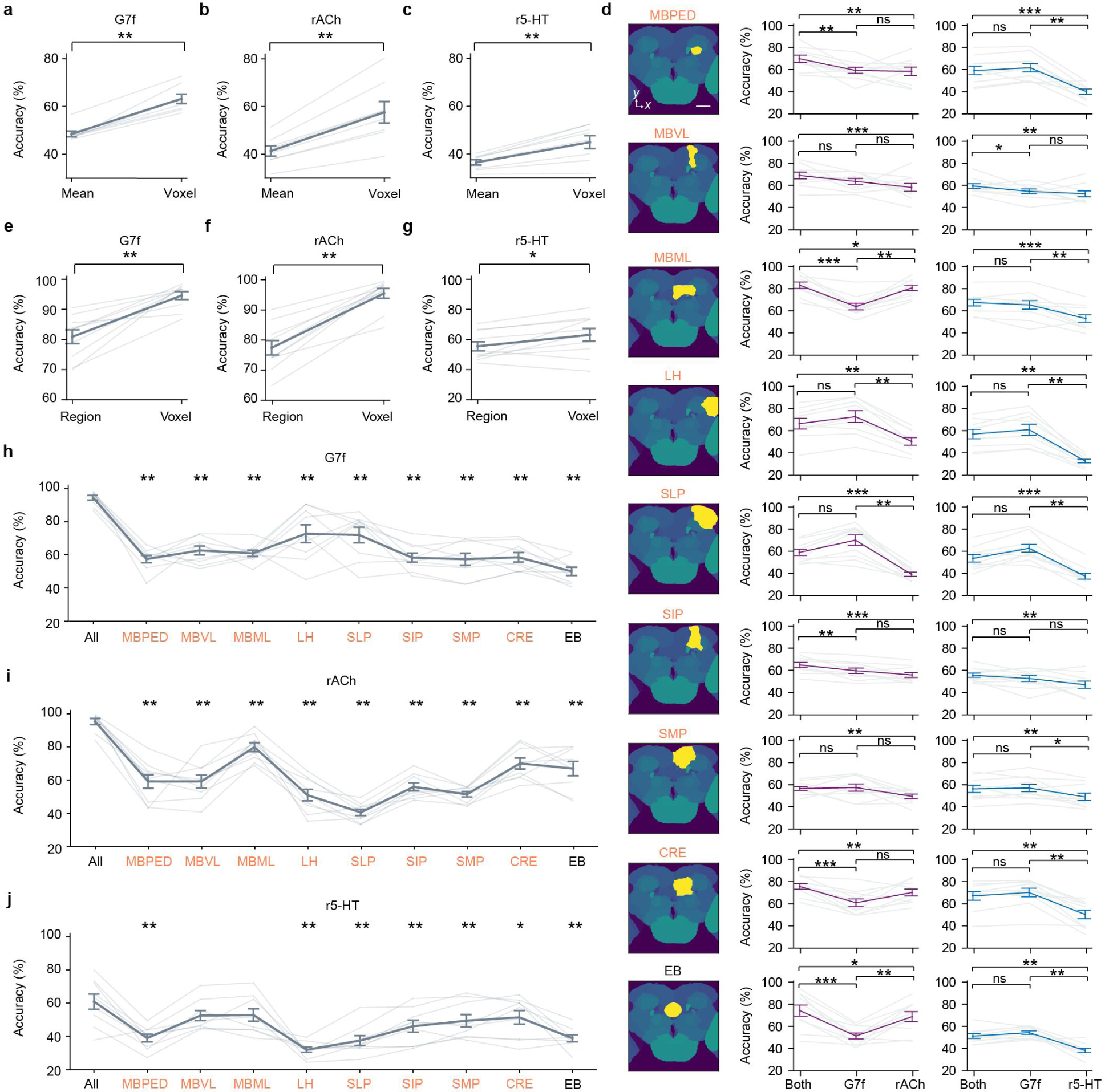
Comparison of odor identity classification accuracies across channels and scales. **a**-**c**, Statistics of the average odor classification accuracy of the blocks and the accuracy integrating all voxels in each olfactory region of the left semi-brain for G7f (**a**), rACh (**b**), and r5-HT (**c**). Each light-colored line represents the average accuracies of a region across all flies. n = 8 regions, mean ± s.e.m. **d**, Comparisons of the voxel-level odor identity classification accuracies by different channels in each brain region (left-side olfactory brain regions and EB). Each region is highlighted in the atlas schematic diagram. **e**-**g**, Statistics of the odor identity classification accuracies of region-level and voxel-level data across multiple brain regions for G7f (**e**), rACh (**f**), and r5-HT (**g**). **h**-**j**, Statistics of the voxel-level odor identity classification accuracies across multiple brain regions and in single brain regions (left-side olfactory brain regions and EB) for G7f (**h**), rACh (**i**), and r5-HT (**j**). Hypothesis testing is performed between “All” and each region. 10 flies co-labeled by G7f and rACh and 10 flies co-labeled by G7f and r5-HT are analyzed. *n* = 10, mean ± s.e.m in **d**-**j**. Results of the G7f channel of the flies co-labeled by G7f and rACh are shown in **a**, **e**, and **h**. Flies without a general coverage of region LH are excluded from the statistics of this region in **a**-**d**. Flies without a general coverage of region LH are excluded for clarity in **h**-**j**. Each light-colored line in the statistical graphs represents the accuracies of a fly in **d**-**j**. Wilcoxon signed-rank test (one-sided test between “Both” and each channel in **d**, and two-sided test for others), \*\*\*\**P* < 0.0001, \*\*\**P* < 0.001, \*\**P* < 0.01, \**P* < 0.05, ns - not significant (*P* > 0.05). Scale bar: 50 μm in **d**.

**Extended Data Fig. 6.**
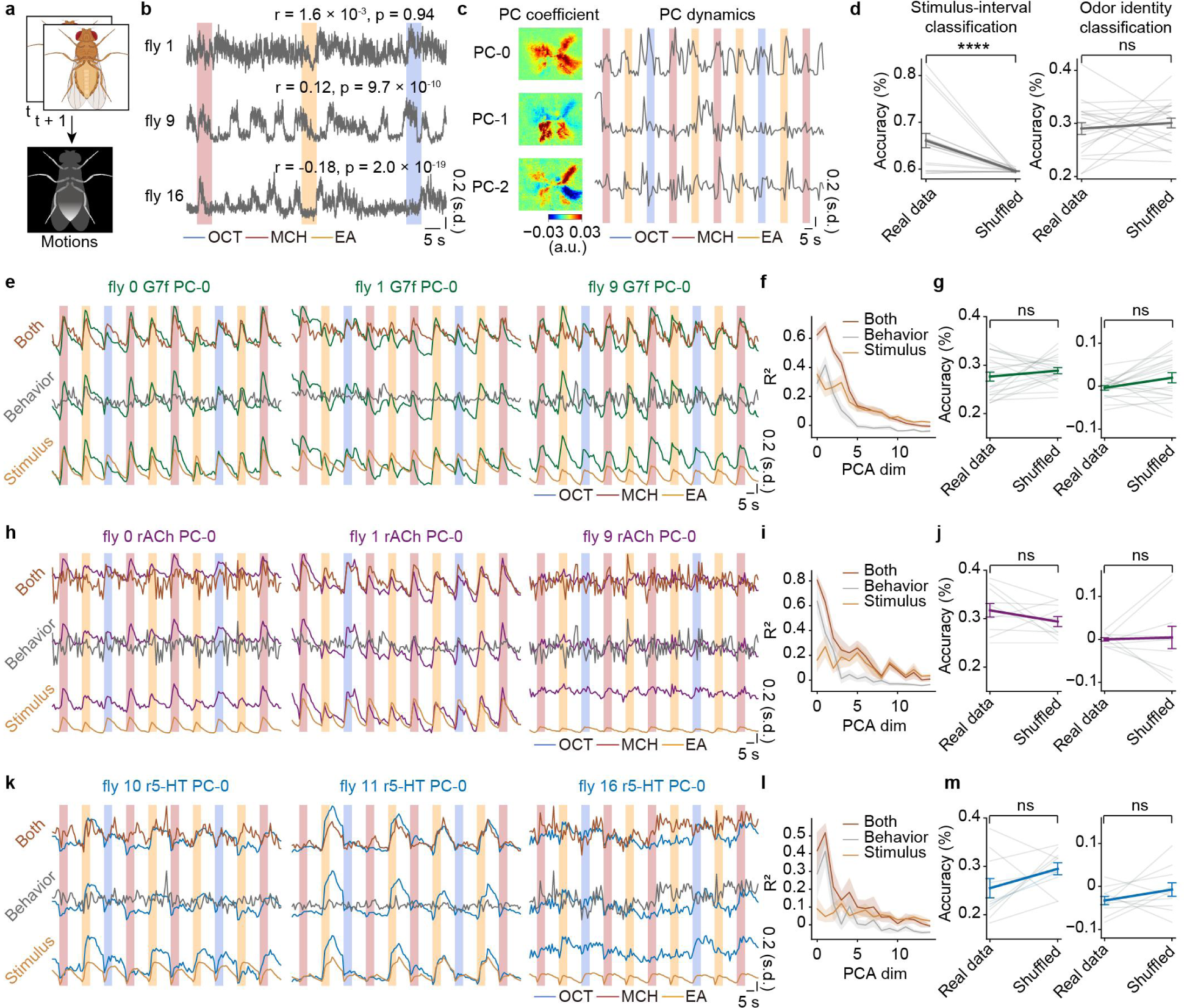
Motion modulates neuronal and neuromodulatory activities but does not account for odor identity representation. **a**, Motions are extracted by subtracting consecutive frames of the videos recording fly abdomens and computing the absolute differences. **b**, Dynamically changing motion energy of the flies. The correlation between odor stimuli and motion energy is diverse and can be weak (top), positive (middle), or negative (bottom). **c**, PCA is performed on the videos to reduce the dimensionality and extract the behavioral features. Left: PC coefficients mapped on the FOV. Right: The dynamics of the PCs in a short period. **d**, Left: The classification accuracy of stimulus periods and intervals. Right: The classification accuracy of odor identities. “Shuffled” refers to shuffling the labels. **e**, Regression of neuronal activities by stimulus (gold, bottom), behavior (gray, middle), and both (brown, top). The green lines are the first PC components of the G7f signals across the brain (step 3 in Fig. 3c). **f**, R^2^ of the regression. The results of the regression of the first 15 PCs are shown. **g**, Left: The classification accuracy of odor identities by the partial neuronal PCs explained by behavior. Right: The change of the classification accuracy of odor identities by removing the neuronal components explained by behavior. “Shuffled” refers to shuffling the labels. **h**-**j**, Similar to **e**-**g**, but showing the regression of the rACh signals. **k**-**m**, Similar to **e**-**g**, but showing the regression of the r5-HT signals. The shades sign odor stimulus periods (blue: OCT; red: MCH; orange: EA) in **b**, **c**, **e**, **h**, and **k**. Each light-colored line represents a fly in **d**, **g**, **j**, and **m**. 10 flies co-labeled by G7f and rACh and 10 flies co-labeled by G7f and r5-HT are analyzed. *n* = 20 flies for G7f and motion analyses, *n* = 10 flies for rACh, *n* = 10 flies for r5-HT, mean ± s.e.m. Two-sided Wilcoxon signed-rank test; \*\*\*\**P* < 0.0001, \*\*\**P* < 0.001, \*\**P* < 0.01, \**P* < 0.05, ns - not significant (*P* > 0.05).

**Extended Data Fig. 7.**
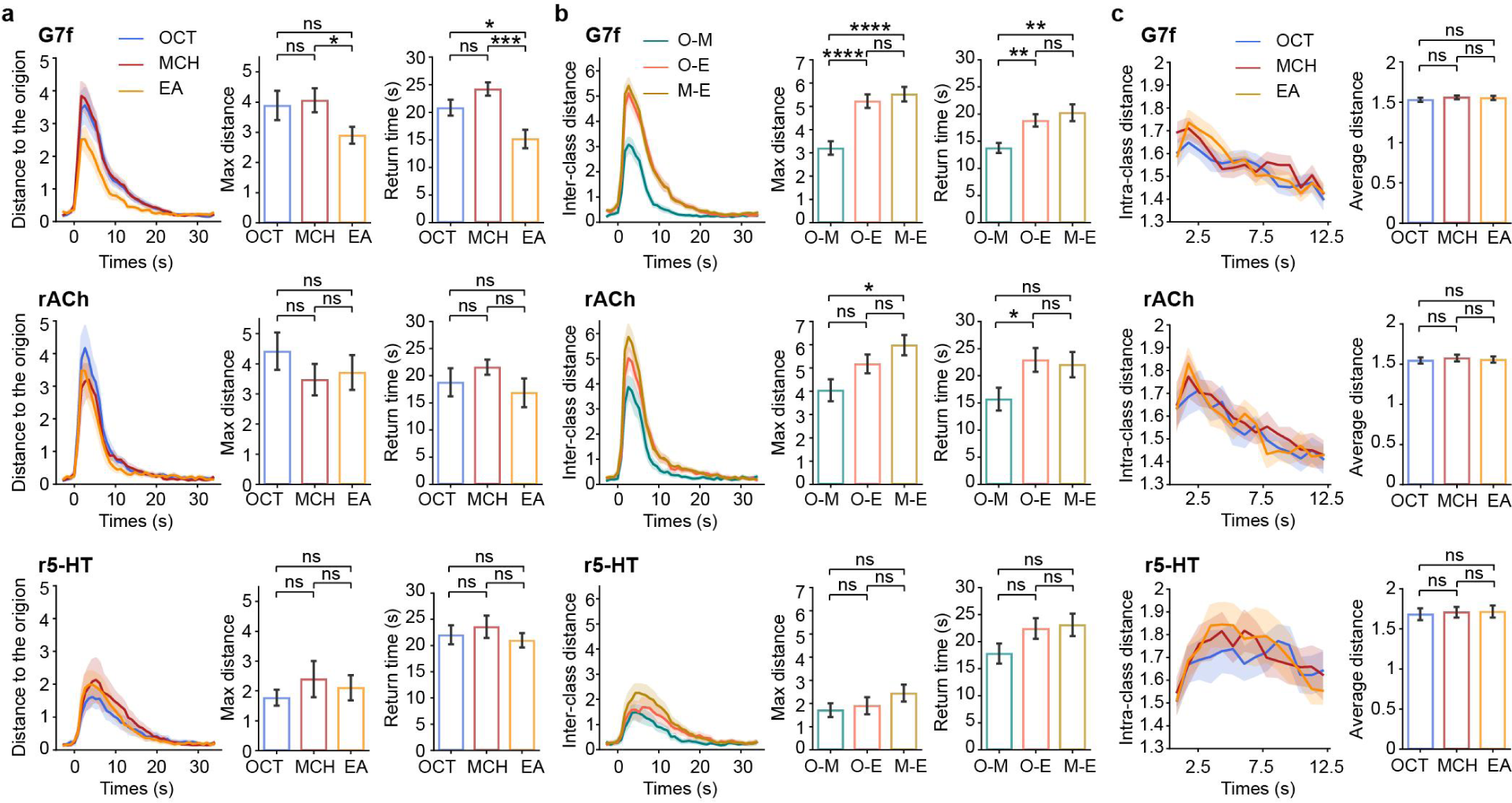
Comparison of the manifolds of different odors. **a**, Comparison of the distance to the origin of different odors. Left: Average distance to the origin changing with time relative to odor delivery. Middle: Statistics of the maximum distance. Right: Statistics of the time for returning to the random state. The three rows show results of G7f, rACh, and r5-HT, from top to bottom, respectively. **b**, Similar to **a**, but for inter-class distance. O-M: OCT and MCH, O-E: OCT and EA, M-E: MCH and EA. **c**, Comparison of the intra-class distance of different odors. Left: Average intra-class distance changing with time relative to odor delivery. Right: Statistics of the average intra-class distance within the period shown in the left figure. 10 flies co-labeled by G7f and rACh and 10 flies co-labeled by G7f and r5-HT are analyzed. *n* = 20 flies for G7f, *n* = 10 flies for rACh, *n* = 10 flies for r5-HT, mean ± s.e.m. Two-sided Mann-Whitney U test; \*\*\*\**P* < 0.0001, \*\*\**P* < 0.001, \*\**P* < 0.01, \**P* < 0.05, ns - not significant (*P* > 0.05).

**Extended Data Fig. 8.**
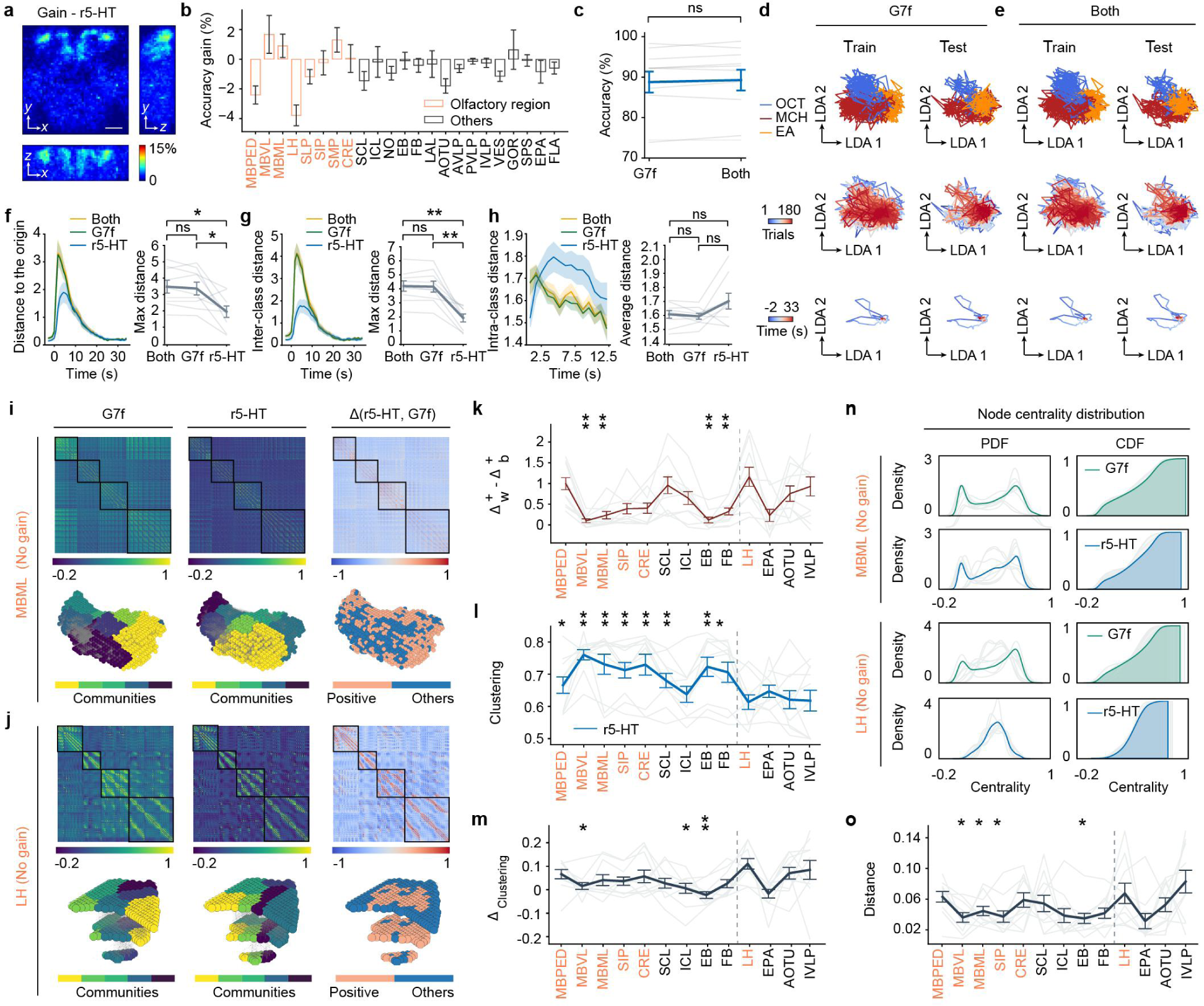
Integration of 5-HT dynamics does not improve the odor identity representation by neuronal activity. **a**, Map of accuracy gain integrating r5-HT dynamics averaged across flies. **b**, Statistics of the average accuracy gain in each brain region. **c**, Comparison of the voxel-level multiple-brain-region odor identity classification accuracies between using only the G7f channel and integrating both channels. **d**, **e**, Low-dimensional manifolds by G7f (**d**) and both channels (**e**) of a fly, similar to Fig. 4d, e. The manifolds are from the same fly as Fig. 3f. The arrow lines are arbitrary units but indicate an equivalent length in each dimension. **f**-**h**, Metrics of the manifolds for integrating both channels, G7f and r5-HT. **f**, Distance to the origin. **g**, Inter-class distance. **h**, Intra-class distance. **i**, **j**, Functional connectivity of the voxels within two brain regions, MBML (**i**) and LH (**j**). Three columns are descriptions of the functional connectivity of G7f, r5-HT, and the difference in the deflation ratio between them, from left to right. For MBML, connectivity emphasis exists without the steady presence of obvious connectivity compensation to G7f by r5-HT (top). The voxels with the greatest increased connectivity do not gather together in space (bottom, coral). For LH, similar results are demonstrated. **k**, The difference in increased deflation ratios within and between clusters of the functional connectivity of r5-HT compared to G7f. **l**, The average clustering coefficient of each brain region for r5-HT. **m**, The average clustering coefficient difference between r5-HT and G7f in each region. **n**, The node centrality distribution of MBML and LH. **o**, The distance of the node centrality distributions between r5-HT and G7f in each region. In **k**, **l**, **m**, and **o**, hypothesis testing is performed as Fig. 4. *n* = 10 flies co-labeled by G7f and r5-HT, mean ± s.e.m. Results of the left-side brain regions and the central complex are shown in **b**, **i**-**o**. Each light-colored line represents the result of a fly in **c**, **f**-**h**, and **k**-**o**. One-sided Wilcoxon signed-rank test in **b**, **l**, **m**, **o**; Two-sided Wilcoxon signed-rank test in **c**, **f-h**, **k**; \*\*\*\**P* < 0.0001, \*\*\**P* < 0.001, \*\**P* < 0.01, \**P* < 0.05, ns - not significant (*P* > 0.05). Scale bar: 50 μm in **a**.

**Extended Data Fig. 9.**
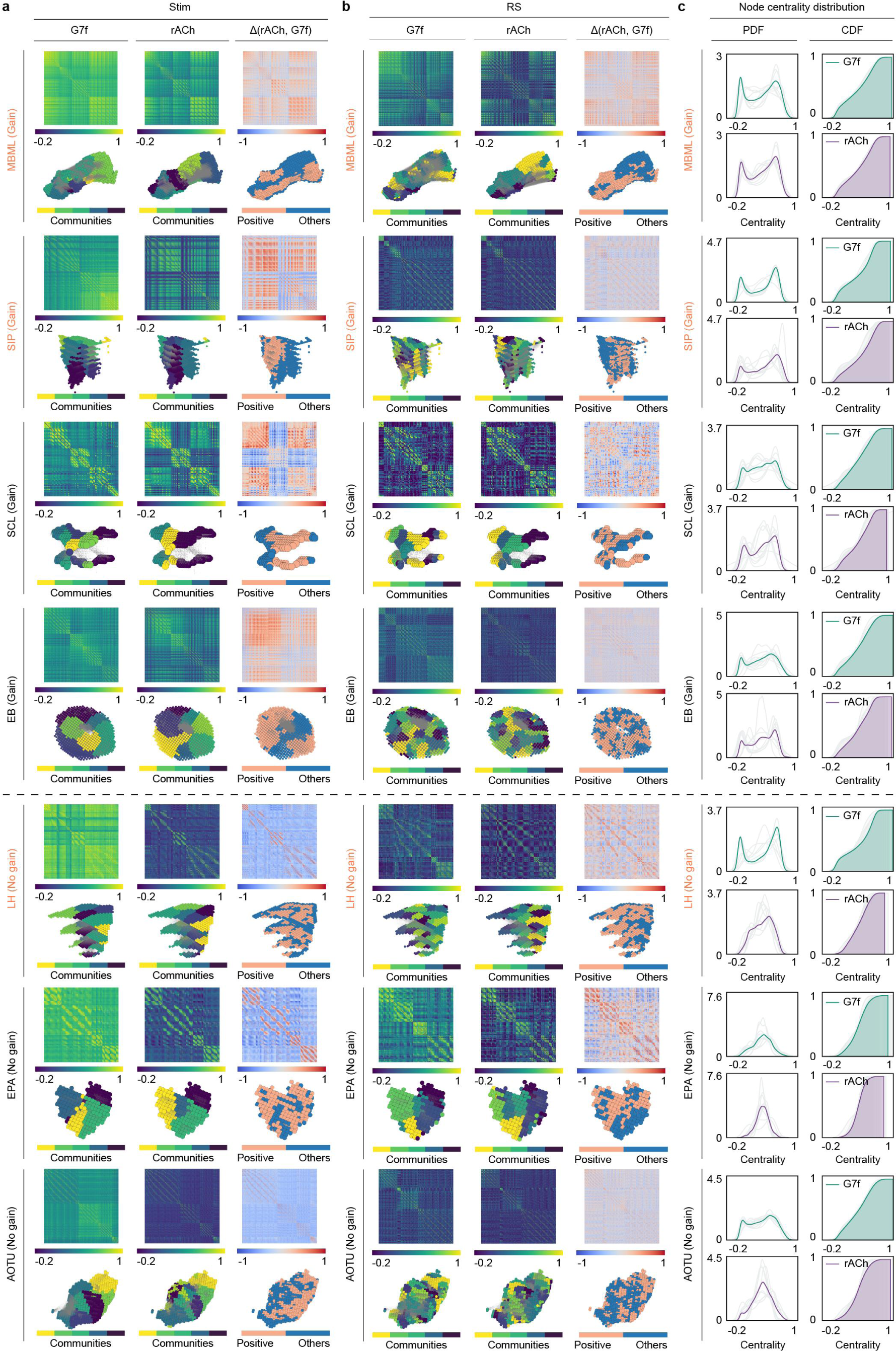
Functional connectivity and topological characterizations by G7f and rACh in brain regions with and without accuracy gain. **a, b**, Functional connectivity matrices and networks by G7f and rACh in four brain regions with accuracy gain and three brain regions without accuracy gain during odor stimulation (**a**) and the resting state (**b**). These seven brain regions are selected from different neuropils of the *Drosophila* brain. For each brain region, three columns are descriptions of the functional connectivity of G7f, rACh, and the difference in the deflation ratio between them, from left to right (top). The functional connectivity of each brain region is shown according to the location of voxels in physical space (bottom). In **a**, for brain regions with accuracy gain, connectivity compensation and emphasis to G7f by rACh are apparent (top). The compensation is reflected in the different distribution of functional connections in the physical space (bottom). The community divisions are similar for the two channels (bottom, labeled in different colors), and voxels with the greatest increased connectivity gather together in space (bottom, coral). Connectivity emphasis exists for brain regions without accuracy gain without obvious compensation and gathering phenomenon. In **b**, the relationship between G7f and rACh is not clearly characterized in the brain regions either with or without accuracy gain. **c**, The node centrality distribution of the brain regions. Brain regions without accuracy gain tend to display more uniform functional connectivity for rACh. Stim: Odor stimulation. RS: The resting state. Each light-colored line represents the result of a fly in **c**.

**Extended Data Fig. 10.**
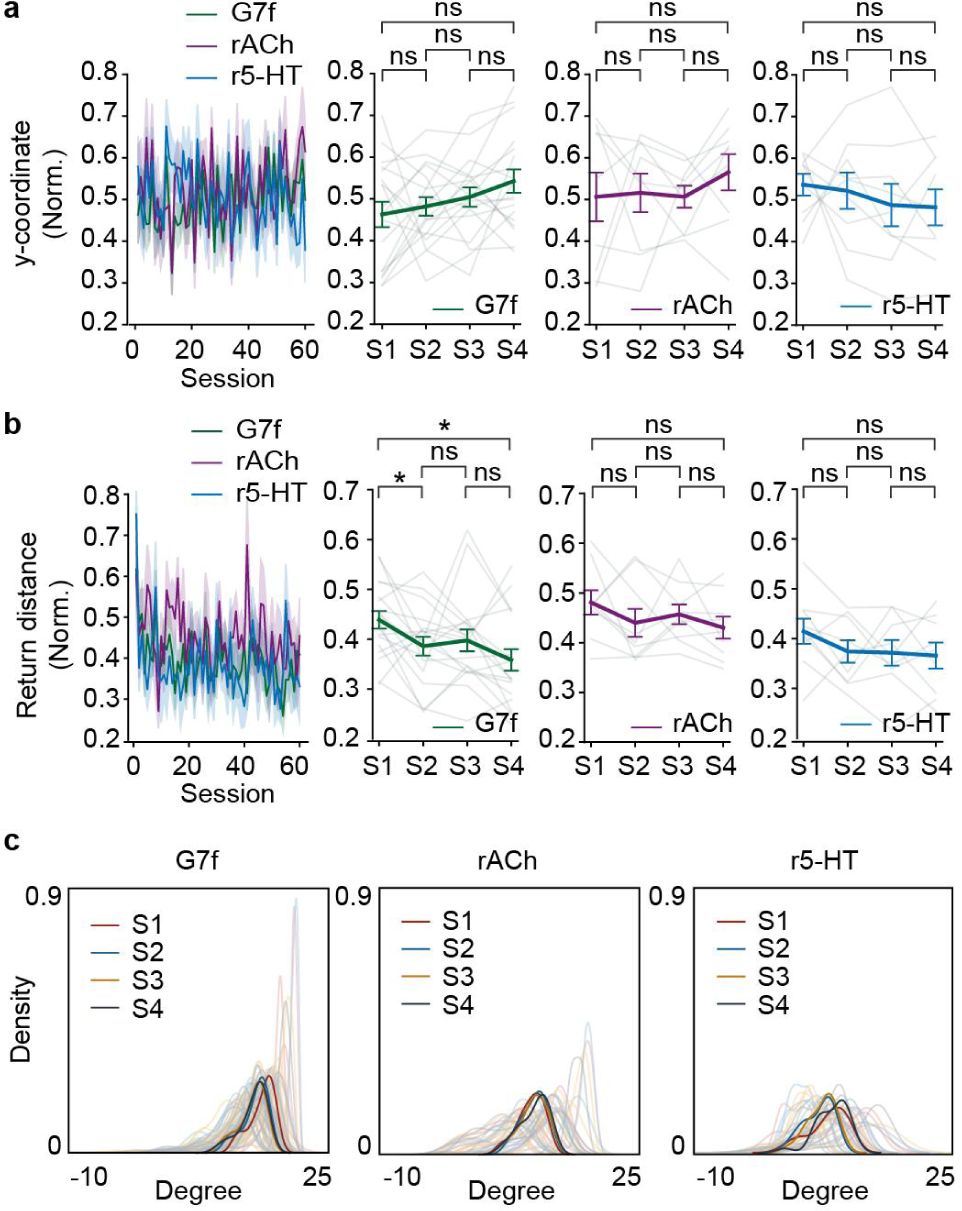
Temporal changes of odor representation and functional connectivity networks. **a**, **b**, Metrics of the manifold changes across four stages. **a**, The y-coordinate of the return locations. **b**, The distance between the return location and the origin. Left: Changes of the metrics across sessions. Right: Statistics of the metrics across four stages for each indicator. **c**, The PDF of the average degree in the functional connectivity networks among four stages (labeled in different colors). Each light-colored line represents a fly. 10 flies co-labeled by G7f and rACh and 10 flies co-labeled by G7f and r5-HT are analyzed. *n* = 20 flies for G7f, *n* = 10 flies for rACh, *n* = 10 flies for r5-HT, mean ± s.e.m. Two-sided Wilcoxon signed-rank test; \*\*\*\**P* < 0.0001, \*\*\**P* < 0.001, \*\**P* < 0.01, \**P* < 0.05, ns - not significant (*P* > 0.05).

**Supplementary Table 1.** The recorded brain regions and the corresponding neuropils. We recorded 43 brain regions in the FOV, including 20 left-side brain regions, 20 right-side brain regions, and 3 brain regions of the central complex. The numbers in the brackets after the neuropil names refer to the number of brain regions recorded. “(L)” indicates that the indices belong to the left-side brain regions, shown in Fig. 2j. Olfactory regions are labeled in coral.

| Neuropil | Brain region index | Brain region name | Full brain region name |
| --- | --- | --- | --- |
| Mushroom body (6) | 64 (L) | MBPED | Pedunculus of mushroom body |
|  | 65 (L) | MBVL | Vertical lobe of mushroom body |
|  | 66 (L) | MBML | Medial lobe of mushroom body |
| Lateral horn (2) | 55 (L) | LH | Lateral horn |
| Superior neuropils (6) | 72 (L) | SLP | Superior lateral protocerebrum |
|  | 73 (L) | SIP | Superior intermediate protocerebrum |
|  | 74 (L) | SMP | Superior medial protocerebrum |
| Inferior neuropils (6) | 63 (L) | CRE | Crepine |
|  | 84 (L) | SCL | Superior clamp |
|  | 59 (L) | ICL | Inferior clamp |
| Central and lateral complex (5) | 4 | NO | Nodulus |
|  | 23 | EB | Ellipsoid body |
|  | 26 | FB | Fan-shaped body |
|  | 56 (L) | LAL | Lateral accessory lobe |
| Ventrolateral neuropils (8) | 79 (L) | AOTU | Anterior optic tubercle |
|  | 75 (L) | AVLP | Anterior ventrolateral protocerebrum |
|  | 76 (L) | PVLP | Posterior ventrolateral protocerebrum |
|  | 77 (L) | IVLP | Wedge |
| Ventromedial neuropils (8) | 60 (L) | VES | Vest |
|  | 80 (L) | GOR | Gorget |
|  | 82 (L) | SPS | Superior posterior slope |
|  | 85 (L) | EPA | Epaulette |
| Periesophageal neuropils (2) | 67 (L) | FLA | Flange |

## Supplementary Video 1

**Volumetric neuronal and neuromodulatory odor responses and low-dimensional manifolds.** This video displays several trials of odor responses of two example flies. The G7f and rACh channels of the fly in Fig. 3d, e, and the r5-HT channel of the fly in Fig. 3f are shown. First row: Volumetric odor responses (mean intensity projections) of G7f, rACh, and r5-HT, from left to right. For the *x*-*z* and *y*-*z* projections, we projected 100 μm where the responses are high along the *y* and *x* directions. Second row: The experimental paradigm, process bar, time and state. The video contains trials 3 - 11 of odor stimulation. Duration from the beginning of the experiment is shown. “Interval” refers to the interval between odor stimulation, and OCT, MCH, EA represent the current stimulus. Third row: The manifolds of this period. The manifolds of single channels and integrating two channels are shown, as signed by the titles of the charts. In each chart, the lines are arbitrary units but indicate an equivalent length in each dimension. Scale bars: 50 μm.

## References

1. Thiele, A. & Bellgrove, M. A. Neuromodulation of Attention. Neuron 97, 769–785 (2018).

2. Lee, S.-H. & Dan, Y. Neuromodulation of Brain States. Neuron 76, 209–222 (2012).

3. Marder, E. Neuromodulation of Neuronal Circuits: Back to the Future. Neuron 76, 1–11 (2012).

4. Xiang, Z., Huguenard, J. R. & Prince, D. A. Cholinergic Switching Within Neocortical Inhibitory Networks. Science 281, 985–988 (1998).

5. Cohn, R., Morantte, I. & Ruta, V. Coordinated and Compartmentalized Neuromodulation Shapes Sensory Processing in *Drosophila*. Cell 163, 1742–1755 (2015).

6. Jacob, S. N. & Nienborg, H. Monoaminergic Neuromodulation of Sensory Processing. Frontiers in Neural Circuits 12, (2018).

7. Linster, C. & Cleland, T. A. Neuromodulation of olfactory transformations. Current Opinion in Neurobiology 40, 170–177 (2016).

8. Lohani, S. et al. Spatiotemporally heterogeneous coordination of cholinergic and neocortical activity. Nat Neurosci 25, 1706–1713 (2022).

9. Thiebaut de Schotten, M. & Forkel, S. J. The emergent properties of the connected brain. Science 378, 505–510 (2022).

10. Lizbinski, K. M. & Dacks, A. M. Intrinsic and Extrinsic Neuromodulation of Olfactory Processing. Frontiers in Cellular Neuroscience 11, (2018).

11. Wilson, D. A., Fletcher, M. L. & Sullivan, R. M. Acetylcholine and Olfactory Perceptual Learning. Learn. Mem. 11, 28–34 (2004).

12. Südhof, T. C. Calcium Control of Neurotransmitter Release. Cold Spring Harb Perspect Biol 4, a011353 (2012).

13. Sturgill, J., et al. Basal Forebrain-Derived Acetylcholine Encodes Valence-Free Reinforcement Prediction Error. http://biorxiv.org/lookup/doi/10.1101/2020.02.17.953141x (2020) doi:10.1101/2020.02.17.953141.

14. Briand, L. A., Gritton, H., Howe, W. M., Young, D. A. & Sarter, M. Modulators in concert for cognition: Modulator interactions in the prefrontal cortex. Progress in Neurobiology 83, 69–91 (2007).

15. Mohebi, A. et al. Dissociable dopamine dynamics for learning and motivation. Nature 570, 65–70 (2019).

16. Juarez, B. et al. Temporal scaling of dopamine neuron firing and dopamine release by distinct ion channels shape behavior. Science Advances 9, eadg8869 (2023).

17. Zhu, F., Elnozahy, S., Lawlor, J. & Kuchibhotla, K. V. The cholinergic basal forebrain provides a parallel channel for state-dependent sensory signaling to auditory cortex. Nat Neurosci 26, 810–819 (2023).

18. Lee, B. K. et al. A principal odor map unifies diverse tasks in olfactory perception. Science 381, 999–1006 (2023).

19. Barnum, G. & Hong, E. J. Olfactory coding. Current Biology 32, R1296–R1301 (2022).

20. Grabe, V. & Sachse, S. Fundamental principles of the olfactory code. Biosystems 164, 94– 101 (2018).

21. Luo, L. Principles of Neurobiology. (Garland Science, 2015).

22. Galizia, C. G. Olfactory coding in the insect brain: data and conjectures. European Journal of Neuroscience 39, 1784–1795 (2014).

23. Pashkovski, S. L. et al. Structure and flexibility in cortical representations of odour space. Nature 583, 253–258 (2020).

24. Campbell, R. A. A. et al. Imaging a Population Code for Odor Identity in the *Drosophila* Mushroom Body. J Neurosci 33, 10568–10581 (2013).

25. Wilson, R. I., Turner, G. C. & Laurent, G. Transformation of Olfactory Representations in the *Drosophila* Antennal Lobe. Science 303, 366–370 (2004).

26. Schoonover, C. E., Ohashi, S. N., Axel, R. & Fink, A. J. P. Representational drift in primary olfactory cortex. Nature 594, 541–546 (2021).

27. Deitch, D., Rubin, A. & Ziv, Y. Representational drift in the mouse visual cortex. Current Biology 31, 4327–4339.e6 (2021).

28. Ogg, M. C., Ross, J. M., Bendahmane, M. & Fletcher, M. L. Olfactory bulb acetylcholine release dishabituates odor responses and reinstates odor investigation. Nat Commun 9, 1868 (2018).

29. Chaudhury, D., Escanilla, O. & Linster, C. Bulbar Acetylcholine Enhances Neural and Perceptual Odor Discrimination. J. Neurosci. 29, 52–60 (2009).

30. Rothermel, M., Carey, R. M., Puche, A., Shipley, M. T. & Wachowiak, M. Cholinergic Inputs from Basal Forebrain Add an Excitatory Bias to Odor Coding in the Olfactory Bulb. J. Neurosci. 34, 4654–4664 (2014).

31. Zhao, Z. et al. Two-photon synthetic aperture microscopy for minimally invasive fast 3D imaging of native subcellular behaviors in deep tissue. Cell 186, 2475–2491.e22 (2023).

32. Dana, H. et al. High-performance calcium sensors for imaging activity in neuronal populations and microcompartments. Nat Methods 16, 649–657 (2019).

33. Jing, M. et al. An optimized acetylcholine sensor for monitoring *in vivo* cholinergic activity. Nat Methods 17, 1139–1146 (2020).

34. Wan, J. et al. A genetically encoded sensor for measuring serotonin dynamics. Nat Neurosci 24, 746–752 (2021).

35. Li, X. et al. Reinforcing neuron extraction and spike inference in calcium imaging using deep self-supervised denoising. Nat Methods 18, 1395–1400 (2021).

36. Li, X. et al. Real-time denoising enables high-sensitivity fluorescence time-lapse imaging beyond the shot-noise limit. Nat Biotechnol 41, 282–292 (2023).

37. Li, X. et al. Spatial redundancy transformer for self-supervised fluorescence image denoising. Nat Comput Sci 3, 1067–1080 (2023).

38. Salvaterra, P. M. & Kitamoto, T. *Drosophila* cholinergic neurons and processes visualized with Gal4/UAS–GFP⋆⋆Published on the World Wide Web on 12 July 2001. Gene Expression Patterns 1, 73–82 (2001).

39. Barnstedt, O. et al. Memory-Relevant Mushroom Body Output Synapses Are Cholinergic. Neuron 89, 1237–1247 (2016).

40. Coletta, L. et al. Network structure of the mouse brain connectome with voxel resolution. Science Advances 6, eabb7187 (2020).

41. Ito, K. et al. A Systematic Nomenclature for the Insect Brain. Neuron 81, 755–765 (2014).

42. Mann, K., Gallen, C. L. & Clandinin, T. R. Whole-Brain Calcium Imaging Reveals an Intrinsic Functional Network in *Drosophila*. Current Biology 27, 2389–2396.e4 (2017).

43. Tully, T., Preat, T., Boynton, S. C. & Vecchio, M. D. Genetic dissection of consolidated memory in *Drosophila*. Cell 79, 35–47 (1994).

44. Shuai, Y., Hu, Y., Qin, H., Campbell, R. A. A. & Zhong, Y. Distinct molecular underpinnings of *Drosophila* olfactory trace conditioning. Proceedings of the National Academy of Sciences 108, 20201–20206 (2011).

45. Schnitzer, M., et al. Dopamine Signals Integrate Innate and Learnt Valences to Regulate Memory Dynamics. https://www.researchsquare.com/article/rs-1915648/v1x (2022) doi:10.21203/rs.3.rs-1915648/v1.

46. Ruebenbauer, A., Schlyter, F., Hansson, B. S., Löfstedt, C. & Larsson, M. C. Genetic Variability and Robustness of Host Odor Preference in *Drosophila* melanogaster. Current Biology 18, 1438–1443 (2008).

47. Stahl, A. et al. Associative learning drives longitudinally graded presynaptic plasticity of neurotransmitter release along axonal compartments. eLife https://elifesciences.org/articles/76712x (2022) doi:10.7554/eLife.76712.

48. Chae, H., Banerjee, A., Dussauze, M. & Albeanu, D. F. Long-range functional loops in the mouse olfactory system and their roles in computing odor identity. Neuron (2022) doi:10.1016/j.neuron.2022.09.005.

49. Schaffer, E. S. et al. The spatial and temporal structure of neural activity across the fly brain. Nat Commun 14, 5572 (2023).

50. Brezovec, L. E., Berger, A. B., Druckmann, S. & Clandinin, T. R. Mapping the Neural Dynamics of Locomotion across the *Drosophila* Brain. 2022.03.20.485047 Preprint at 10.1101/2022.03.20.485047 (2022).

51. Aimon, S. et al. Fast near-whole–brain imaging in adult *Drosophila* during responses to stimuli and behavior. PLOS Biology 17, e2006732 (2019).

52. Hige, T., Aso, Y., Rubin, G. M. & Turner, G. C. Plasticity-driven individualization of olfactory coding in mushroom body output neurons. Nature 526, 258–262 (2015).

53. Chae, H. et al. Mosaic representations of odors in the input and output layers of the mouse olfactory bulb. Nat Neurosci 22, 1306–1317 (2019).

54. Allard, A., Serrano, M. Á. & Boguñá, M. Geometric description of clustering in directed networks. Nat. Phys. 1–7 (2023) doi:10.1038/s41567-023-02246-6.

55. Steinmetz, N. A., Zatka-Haas, P., Carandini, M. & Harris, K. D. Distributed coding of choice, action and engagement across the mouse brain. Nature 576, 266–273 (2019).

56. Ebrahimi, S. et al. Emergent reliability in sensory cortical coding and inter-area communication. Nature 605, 713–721 (2022).

57. Pacheco, D. A., Thiberge, S. Y., Pnevmatikakis, E. & Murthy, M. Auditory activity is diverse and widespread throughout the central brain of *Drosophila*. Nat Neurosci 24, 93–104 (2021).

58. Allen, W. E. et al. Thirst regulates motivated behavior through modulation of brainwide neural population dynamics. Science 364, eaav3932 (2019).

59. Wandell, B. A., Dumoulin, S. O. & Brewer, A. A. Visual field maps in human cortex. Neuron 56, 366–383 (2007).

60. Kannan, M. et al. Dual-polarity voltage imaging of the concurrent dynamics of multiple neuron types. Science 378, eabm8797 (2022).

61. Mu, Y. et al. Glia Accumulate Evidence that Actions Are Futile and Suppress Unsuccessful Behavior. Cell 178, 27–43.e19 (2019).

62. Deng, F. et al. Dual-color GRAB sensors for monitoring spatiotemporal serotonin release *in vivo*. 2023.05.27.542566 Preprint at 10.1101/2023.05.27.542566 (2023).

63. Ji, N., Freeman, J. & Smith, S. L. Technologies for imaging neural activity in large volumes. Nat Neurosci 19, 1154–1164 (2016).

64. Stringer, C., Michaelos, M., Tsyboulski, D., Lindo, S. E. & Pachitariu, M. High-precision coding in visual cortex. Cell 184, 2767–2778.e15 (2021).

65. Pinto, L. et al. Fast modulation of visual perception by basal forebrain cholinergic neurons. Nat Neurosci 16, 1857–1863 (2013).

66. Li, T. et al. Cellular bases of olfactory circuit assembly revealed by systematic time-lapse imaging. Cell 184, 5107–5121.e14 (2021).

67. Xiao, N. et al. A single photoreceptor splits perception and entrainment by cotransmission. Nature 1–9 (2023) doi:10.1038/s41586-023-06681-6.

68. Chantranupong, L. et al. Dopamine and glutamate regulate striatal acetylcholine in decision-making. Nature 621, 577–585 (2023).

69. Mann, K., Deny, S., Ganguli, S. & Clandinin, T. R. Coupling of activity, metabolism and behaviour across the *Drosophila* brain. Nature 593, 244–248 (2021).

70. Yang, Q. et al. Spontaneous recovery of reward memory through active forgetting of extinction memory. Current Biology 33, 838–848.e3 (2023).

71. Mo, H. et al. Age-related memory vulnerability to interfering stimuli is caused by gradual loss of MAPK-dependent protection in *Drosophila*. Aging Cell 21, e13628 (2022).

72. Murthy, M. & Turner, G. Dissection of the Head Cuticle and Sheath of Living Flies for Whole-Cell Patch-Clamp Recordings in the Brain. Cold Spring Harb Protoc 2013, pdb.prot071696 (2013).

73. Pnevmatikakis, E. A. & Giovannucci, A. NoRMCorre: An online algorithm for piecewise rigid motion correction of calcium imaging data. Journal of Neuroscience Methods 291, 83– 94 (2017).

74. Wang, Z. et al. Real-time volumetric reconstruction of biological dynamics with light-field microscopy and deep learning. Nat Methods 18, 551–556 (2021).

75. Kantorovich, L. V. Mathematical Methods of Organizing and Planning Production. Management Science 6, 366–422 (1960).

76. Cramér, H. On the composition of elementary errors. Scandinavian Actuarial Journal 1928, 13–74 (1928).

